# Stress granules formed during different RNA virus infections show remarkable plasticity and substantial virus-specific differences in their formation and composition

**DOI:** 10.1101/2025.08.14.670263

**Authors:** Reto M. Lang, Silvio Steiner, Jenna Kelly, Anne-Christine Uldry, Pratik Dave, Sophie Braga-Lagache, Jeffrey Chao, Manfred Heller, Volker Thiel

**Affiliations:** Institute of Virology and Immunology (IVI), Bern and Mittelhäusern, Switzerland; Department of Infectious Diseases and Pathobiology (DIP), Vetsuisse Faculty, University of Bern, Bern, Switzerland; Graduate School for Cellular and Biomedical Sciences, University of Bern, Bern, Switzerland; Multidisciplinary Center for infectious Diseases (MCID), University of Bern, Bern, Switzerland; Department for BioMedical Research (DBMR), University of Bern, Bern, Switzerland; Friedrich Miescher Institute for Biomedical Research (FMI), Basel, Switzerland

## Abstract

Eukaryotic cells evolved a cellular stress response to cope with extrinsic and intrinsic stress stimuli including virus infections. The major result of this response is the shutdown of bulk translation to prevent damage and allow the reprogramming of translation towards stress-resolving pathways. The resulting translationally stalled mRNA and associated proteins are accumulated in membrane-less cytosolic condensates called stress granules (SG). While the inhibitory effect of translation arrest on virus growth is well established, the role of SGs in the cellular defense against viruses is still unclear. The observation of specific interference with SG formation during various virus infections led to the hypothesis that SGs could serve as antiviral signaling platforms.

In this study, we used mouse hepatitis virus (MHV) to characterize SGs formed during coronavirus infection. By applying APEX2-mediated proximity labelling in combination with quantitative proteomics, we dissected the proteome of MHV-induced granules and compared it to canonical SGs formed during oxidative stress. Our data revealed substantial differences in protein abundance and composition indicating stressor-specific SG characteristics. To assess if the observed differences are a general feature of virus-induced SGs or rather virus-specific, we extended our investigations to the Semliki Forest virus (SFV), a member of the alphavirus family known to induce SGs. An initial comparison of SG formation kinetics by live-cell imaging showed distinct time points of SG induction between both viruses. A comprehensive comparison of the SG protein compositions revealed profound differences in the SG proteome between SFV and MHV. A further subcellular localization of SG components by microscopy not only confirmed a reduced abundance of several translation initiation factors in MHV-induced granules, but surprisingly, revealed the presence of SFV RNA and the absence of MHV RNA in virus-induce SGs.

The reduced connection to canonical SG themes observed for MHV-induced granules compared to SFV- and oxidative stress-induced ones indicates a different impact of these condensates on MHV replication and further raises the question whether they should be considered SGs. The surprising plasticity of SGs concerning induction kinetics, protein composition and abundance, and inclusion or exclusion of viral RNA provide a base for future investigations of the role(s) of SGs in the context of viral infection and how they may impact virus replication.

## Introduction

Eukaryotic cells have evolved a stress response to cope with various intrinsic and extrinsic stress stimuli including virus infections. Besides a shut-down of bulk translation, this response is characterized by the formation of cytosolic stress granules (SGs). These highly dynamic condensates consist of translationally stalled mRNAs and various associated proteins including eukaryotic translation initiation factors (eIFs) (1,2). The formation of SGs is mediated by specific RNA-binding proteins like the major SG markers Ras GTPase-activating protein-binding protein 1 (G3BP1), T-cell restricted intracellular antigen-1 (TIA-1) and ubiquitin associated protein 2 like (UBAP2L) by promoting a multivalent network of RNA and protein interactions (1,3,4). Many studies have suggested the involvement of SGs in the antiviral response as a signaling platform. Several antiviral sensor proteins including the protein kinase R (PKR) were found to accumulate in SGs during different stress stimuli (5–7). Furthermore, various viruses have been shown to interfere with the SG response by inhibiting their formation or manipulating their characteristics to favor viral replication (8,9). However, the proposed function of SGs as an antiviral signaling platform was recently questioned, as several studies could not find antiviral sensor proteins in SGs and further the absence of SGs by G3BP1-KO did not alter antiviral signaling (10,11). These contradictory results show that the role of SGs during virus infections is still poorly understood and might vary between viruses.

Aside from mRNAs, SGs accumulate a large variety of cellular proteins, which might modulate their function during different stress stimuli. To comprehensively identify the proteome of these highly dynamic condensates and capture transiently interacting proteins two studies previously used proximity labelling assays based on the biotin ligase BirA or the engineered peroxidase APEX2 (12,13). APEX2 is fused to G3BP1 as SG bait and catalyzes the tagging of G3BP1 proximal proteins with biotin phenol (BP) during a short labelling period of one minute (12). Markmiller and colleagues applied APEX2-mediated proximity labelling in combination with quantitative mass spectrometry to compare the SG proteome during different non-viral stress situations including oxidative stress induced by sodium arsenite treatment (12). Among the classical SG marker proteins, they identified many novel SG associated proteins. Furthermore, Markmiller et al. and others found that the protein composition of SGs differs between stress stimuli as well as cell types (1,12,13). These observations demonstrate stress type specific characteristics of SGs suggesting an adaptation of these condensates to handle different stress conditions.

As the characteristics of SGs in the context of virus infections are still poorly understood, we set out to compare SGs induced by two different viruses, the Semliki Forest virus (SFV) and the mouse hepatitis virus (MHV). While many viruses prevent SG formation completely, the Semliki Forest virus induces SGs early in infection by activating the dsRNA sensor PKR (9,14). This virus is a member of the alphavirus family possessing a positive-sense single stranded, capped and polyadenylated RNA genome. Interestingly, SFV was shown to actively dissolve the granules later in infection and prevent their reformation (9,15). The mechanism was suggested to be mediated by the interaction of nsP3 with G3BP1 and the subsequent sequestration of this SG core component (9,16). However, a recent study questions this view by showing that a loss of the nsp3-G3BP1 interaction does not prevent SG disassembly, while SFV-induced transcription and translation shut off seem to be essential (17). Nevertheless, the formation of SG in infection and their subsequent dissolvement by the virus, makes SFV a suitable model to study these granules in virus infections.

MHV is a member of the coronavirus family, which is characterized by an exceptionally long positive-sense single stranded RNA genome. Coronaviruses encode various non-structural and accessory proteins, which are involved in the evasion of host defense mechanisms (18). In the context of the stress response, it was recently found that the human coronavirus SARS-CoV-2, causative agent of the COVID19 pandemic, inhibit SG formation mediated by the interaction of the nucleocapsid protein (N) with G3BP1 (10,19). Interestingly, this interaction motif in N is specific for SARS-CoV-2 and could not been found in other coronaviruses including MHV, which was previously shown to trigger SG formation during infection (20). However, the role of these SGs during coronavirus infections as well as the general characteristics of virus-induced SGs is still poorly understood. Due to a possible role in the antiviral response, the insight into SGs induced by different viruses can provide valuable information on the host-virus interaction in the context of the stress response. This is of special importance for coronaviruses, as the recent COVID-19 pandemic demonstrated the risk posed by members of this virus family and the need to understand their interaction with the host cell.

Here we provide a detailed characterization of SGs in the context of virus infections. By comparing these condensates induced by two RNA viruses from the coronavirus and alphavirus families we observed substantial differences in the kinetics of SG formation. By applying APEX2-mediated proximity labelling, quantitative proteomics and different microscopy-based techniques, we report a remarkable plasticity of SGs concerning protein composition and abundance and concerning the inclusion or exclusion of viral RNA. Collectively, our data on various aspects of virus-induced SGs unravels virus-specific characteristics of these condensates that may impact their role(s) during virus infection.

## Results

### Generation of G3BP1-APEX2-GFP expressing cell line for microscopy-based investigations and proximity labelling

To investigate SGs formed during infection with MHV, we first generated a recombinant mouse fibroblast 17Cl1 cell line stably expressing a fusion protein of the SG marker G3BP1, the engineered ascorbate peroxidase APEX2, and an eGFP tag, hereafter referred to as G3AG (Fig 1A). While the eGFP enables microscopy-based characterization of SG formation, the APEX2 enzyme is used for proximity-dependent biotinylation allowing dissection of the G3BP1 microenvironment

**Fig 1.**
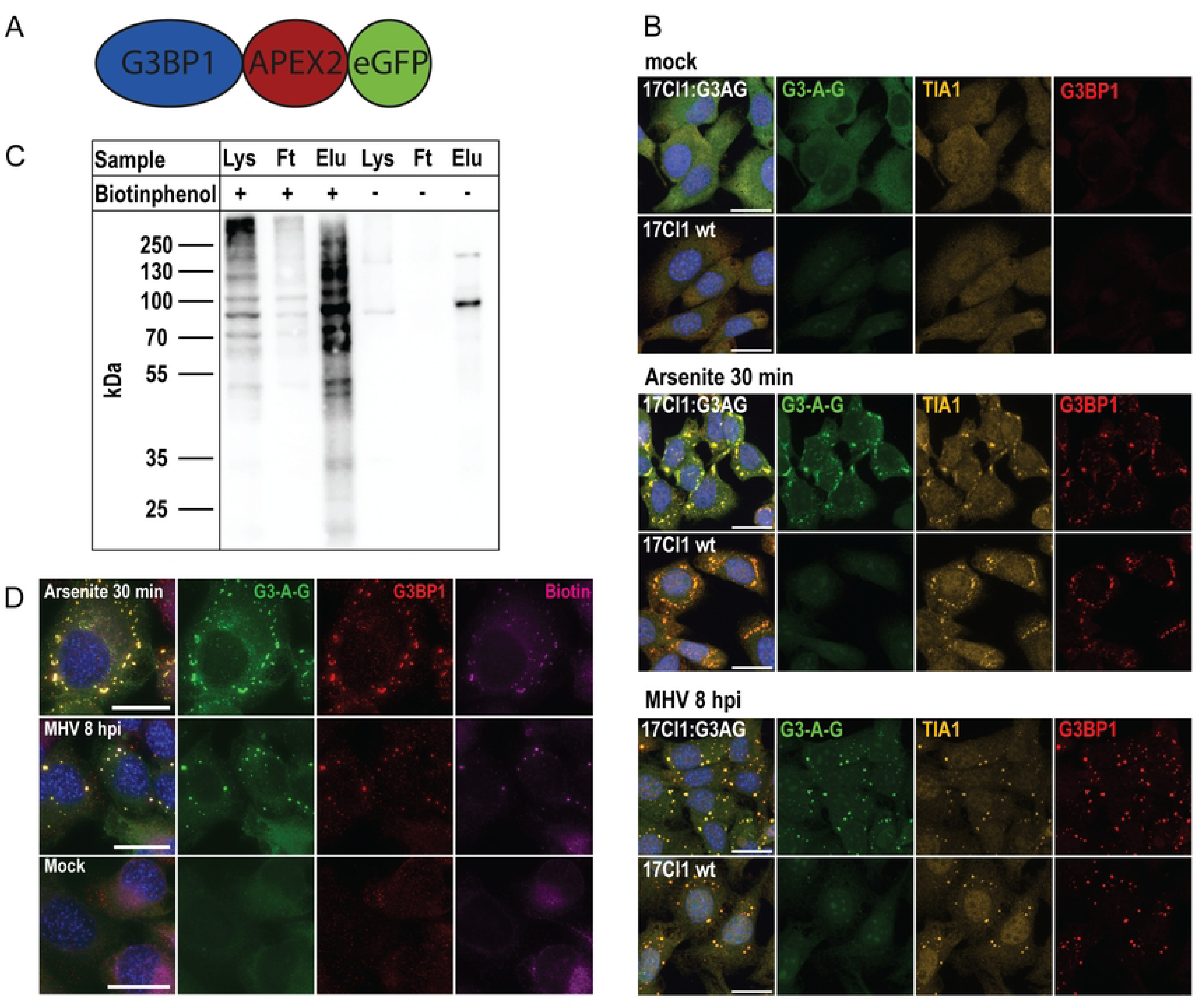
Cell line expressing G3BP1-APEX2-GFP fusion protein allows visualization and proximity-dependent biotinylation of SGs during oxidative and infection stress. (A) Schematic representation of the fusion protein consisting of the main SG marker G3BP1, the engineered APEX2 peroxidase for proximity labelling, and a GFP tag for visualization of SGs by fluorescence microscopy. (B) Verification of SG formation and accumulation of G3-A-G fusion protein (green) in recombinant 17Cl1:G3-A-G cells compared to wildtype cells by immunofluorescence microscopy. 17Cl1:G3-A-G or wildtype cells were treated with 250uM sodium arsenite for 30min or infected with MHV-A59 (MOI 10). Control of unstressed (mock) cells was additionally included. Cells were fixed at the indicated time point after treatment/infection and an indirect immunofluorescence staining targeting TIA1 (orange) and G3BP1 (red) was performed. Nuclei were stained with DAPI (blue). Scale bar: 20um (C) Verification of APEX2-mediated proximity labelling of proteins by Western blot analysis. 17Cl1:G3-A-G were incubated with 500uM biotin phenol (BP) for 30min. Then, proximity labelling reaction was started by adding hydrogen peroxide. After one minute the reaction was quenched and cells were lysed. Affinity purification of labeled proteins was performed on streptavidin-coated magnetic beads. Lysates (Lys), flowthrough (Ft) and eluates (Elu) were analyzed for the presence of biotinylated proteins by Western blot analysis. As negative controls, BP was omitted for the labelling reaction. (D) Verification of APEX2-mediated proximity labelling of SG components by immunofluorescence. Recombinant 17Cl1:G3-A-G cells (green) were either treated with sodium arsenite (250uM, 30min) or infected with MHV-A59 (MOI 5, 8hpi) to induce G3-A-G granules. Unstressed cells (mock) were used as negative control. Syncytia formation in MHV infected cells was prevented by HR2 peptide addition at 2hpi. Cells were incubated with BP 30min before the labeling time points. At the indicated time points, labeling reaction was started by adding hydrogen peroxide. Reaction was quenched after 1min and cells were fixed. An immunofluorescence staining with NeutrAvidin (magenta) to visualize biotinylated proteins, and an antibody targeting the SG marker G3BP1 (red) was performed. Scale bars: 20um

First, we confirmed the accumulation of fusion protein in SGs upon different stress stimuli. We treated 17Cl1:G3AG cells with sodium arsenite, a known inducer of oxidative stress (21), or infected them with MHV strain A-59. The subcellular distribution of fusion protein during presence of SGs was assessed by fluorescence microscopy. Based on a preliminary experiment with wildtype cells, we selected 8hpi as time point of interest for MHV-induced SGs (S1 Fig). We used antibody staining for SG markers G3BP1 and TIA1 to visualize SGs (1). In the unstressed state (mock) the fusion protein is diffusely distributed in the cytosol as visible in figure 1B (top panel). Upon stress induced by either arsenite treatment or MHV infection, G3AG accumulates in cytosolic foci, which overlap with the staining for G3BP1 and TIA1 (Fig 1B, middle panel). These accumulations have a similar appearance to the G3BP1/TIA1 positive SGs formed in the wildtype cells. Based on these results we conclude that G3AG localizes to SGs during stress.

Proximity dependent labelling based on the engineered peroxidase APEX2 has proven an important method to comprehensively identify the proteome of cellular organelles, like mitochondria, as well as more dynamic subcellular structures including SGs (12). To verify the activity of APEX2, we performed a proximity labelling assay in 17Cl1:G3AG cells with a 1min labelling time after 30min of incubation with or without biotin phenol (BP) and further performed affinity purification of labeled proteins. The presence of biotinylated proteins in the lysates, flowthrough and eluates was assessed by Western blot analysis. The lysate of 17Cl1:G3AG cells incubated with BP contains many different biotinylated proteins as indicated by the numerous distinct bands and the diffuse staining distributed over the whole mass range (Fig 1C, first lane). The affinity purification of this sample shows a successful enrichment of labeled proteins in the eluate (Fig 1C, 3^rd^ lane). In contrast, lysate omitted with BP shows only two distinct bands and no diffuse staining (Fig 1C, 4^th^ lane). Taken together, these results verify APEX2-mediated biotinylation of proteins in our fusion protein expressing cells.

We further analyzed the subcellular localization of biotinylated proteins by fluorescence microscopy to verify the labelling of SG components. MHV-infected or sodium arsenite treated 17Cl1:G3AG cells were proximity labelled and fixed at 8hpi and 30min treatment respectively and stained with fluorophore-conjugated neutravidin. The results revealed a strong accumulation of biotinylated proteins in MHV- and arsenite induced SGs, verifying the labelling of SGs (Fig 1D).

Taken together, our data showed that the G3AG fusion protein localizes to SGs upon different stress stimuli. Furthermore, the APEX2 enzyme contained in the fusion protein efficiently catalyzes the proximity-dependent biotinylation of SG proteins.

### MHV-induced stress granules reveal a distinct proteome from canonical stress granules

After verifying the 17Cl1:G3AG cells, we applied APEX2-mediated proximity labelling to comprehensively dissect the proteome of MHV-induced SGs and compare it to canonical SGs induced by sodium arsenite. We combined this technique with stable isotope labelling of amino acids in cell culture (SILAC) allowing a direct comparison of two conditions by quantitative mass spectrometry based on ‘heavy’ or ‘light’ stable isotope labeled amino acids (lysine and arginine) in the culture medium (S2A Fig). We designed five individual comparisons, each of which consists of two different conditions, whereby one is carried out on ‘heavy’ cells and one on ‘light’ cells (S2B Fig). Our setup consists of a direct comparison of MHV- and arsenite-induced SGs as well as a comparison of each stressor with the unstressed condition (Fig 2A/B/C) and a stressed but unlabeled control. To capture the SG proteome, the labelling reaction was performed at 8hpi for MHV infection and 30min after sodium arsenite treatment, respectively. A total of 2206 different proteins were detected over all analyzed samples, including 194 SG associated proteins based on an assembled list of known SG proteins from previous studies (12) and the MGI database.

**Fig 2.**
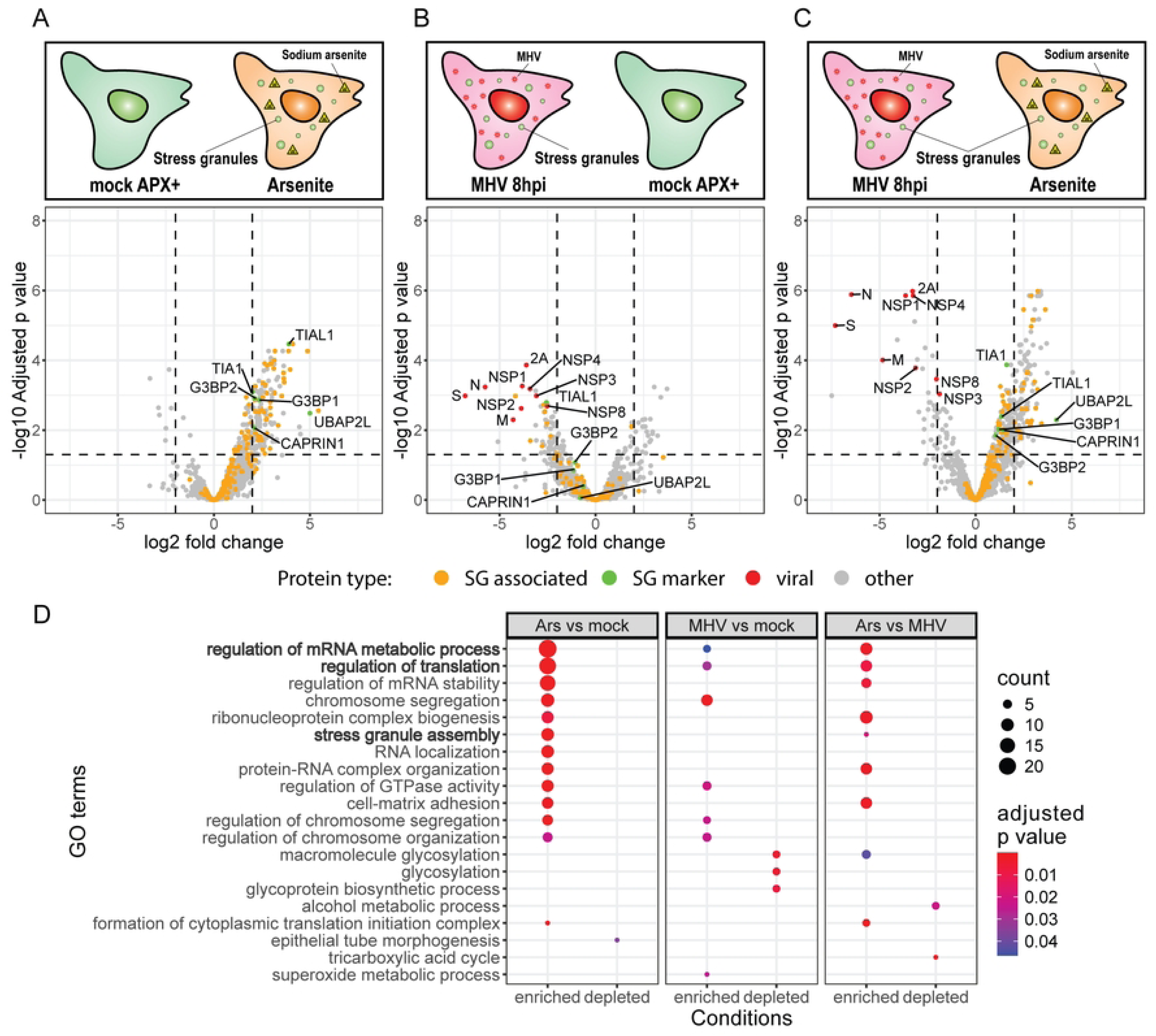
Comparing the protein composition of the SG microenvironment during oxidative stress and MHV infection using APEX2-mediated proximity labelling and quantitative proteomics. Protein composition of the G3-A-G microenvironment was compared between (A) sodium arsenite treated and unstressed cells, (B) MHV-infected and unstressed cells, as well as (C) a direct comparison of the two stressors, sodium arsenite treatment and MHV infection. Volcano plots show the mean log2 fold change between the compared conditions (shown in scheme) for all identified proteins against the corresponding – log10 adjusted p value. Proteins from our SG associated proteins reference list are indicated in orange, main SG marker proteins are labeled and indicated in green, and viral proteins are labeled and indicated in red. The selected significance levels for log2 fold change (≥2 and ≤-2) and adjusted p value (<0.05) are shown with dashed lines. (D) GO term enrichment analysis for biological processes was performed on the fraction of significantly enriched/depleted proteins for each comparison using ClusterProfiler in RStudio. Redundant terms were reduced by semantic similarity using the rrvigo package. The circle size indicates the number of genes identified, the color gradient indicates the adjusted p value (based on Benjamin-Hochberg). Top 5 terms for each condition based on the adjusted p value are displayed. Interesting terms in the context of SGs are highlighted in bold.

We analyzed the data using a similar approach to Markmiller et al. (12). An initial analysis of heavy/light ratios revealed substantial differences in the proteome of MHV- and arsenite-induced SGs. The second analysis based on individual samples, additionally considering proteins unique for one condition, further strengthened this observation. The comparison of arsenite induced SGs with the unstressed state (mock) shows a shift of many SG associated proteins towards the arsenite condition as seen in volcano plot 2A. Among the 147 significantly enriched proteins for arsenite stress based on our stringent significance criteria (log2(fold change) >1), we identify 46 SG associated proteins. These include the well-characterized SG nucleators G3BP1, G3BP2 (22,23), TIA1, TIAL (1,24), CAPRIN1 (23,25) and UBAP2L (4,12,23), as well as several components of the translation initiation complex (Fig 3A). Additionally, our screen identified 69 proteins enriched in arsenite-induced SGs which are not contained in our SG reference list and thus might represent novel candidates of oxidative stress induced SGs. Interestingly, we detected 144 known SG associated proteins, which did not meet our significance criteria. Gene ontology enrichment analysis for biological processes of the significantly enriched proteins showed an enrichment for translation initiation and SG assembly, supporting the successful labelling of the SG microenvironment (Fig 2D). Overall, this experiment shows a strong stress induction by sodium arsenite treatment leading to a change in G3BP1 microenvironment in line with canonical SG formation.

**Fig 3.**
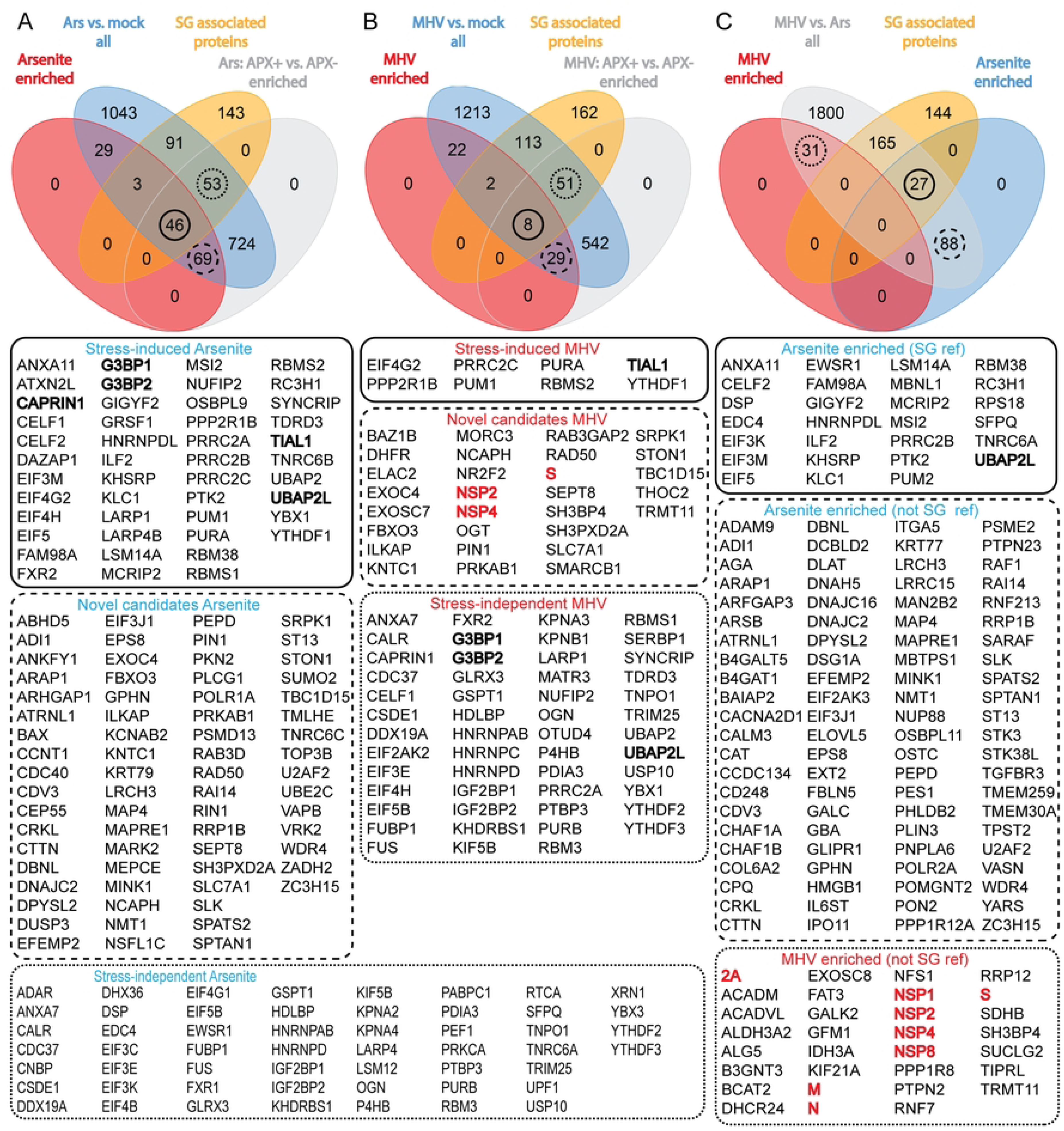
Comparison of SG proteome during oxidative stress and MHV infection. Comparison of SG microenvironment between (A) oxidative stress induced by sodium arsenite treatment and the unstressed state (mock) or (B) MHV infection and the unstressed state (mock). Different subsets of stress-dependent and independent SG proteins are determined using a Venn diagram between the subset of significantly enriched proteins for the stressor (Ars/MHV enriched, red), all identified proteins of the same comparison (blue), the subset of significantly enriched proteins for the proximity labelled condition (APX+, gray) in the negative labelling control comparison, and a reference list of known SG associated proteins (orange). Proteins of specific subsets, including stress induced known SG proteins (solid line), novel candidates of SG proteins for respective stress stimulus (large dashes) and stress independent known SG proteins (small dashes), are highlighted with circles and corresponding lists of proteins are provided below. (C) Venn diagram shows overlap between significantly enriched proteins for arsenite treatment (blue), significantly enriched proteins for MHV infection (red) and all identified proteins in the direct comparison between these stressors (gray) with the SG associated proteins reference list (orange). Subsets of arsenite specific SG proteins (solid line), arsenite specific novel SG protein candidates (large dashes) and MHV specific novel SG candidates (small dashes) are highlighted with circles and corresponding protein lists are provided below. Main SG marker proteins are highlighted in bold, viral proteins in bold and red. For all significantly enriched fractions log2 fold change significance level of ≥2 and ≤-2, and an adjusted p value significance level of <0.05 were used.

In the comparison of the G3BP1 microenvironment between MHV-induced SGs and the unstressed mock cells, we observed the significant enrichment of 61 proteins for MHV. Interestingly, only 10 known SG associated proteins have been detected among these hits with TIAL1 as only SG nucleator protein (Figs 2B/3B). Although additional 164 SG associated proteins have been detected, they show no significant change in abundance during MHV infection. Furthermore, although the GO terms “Regulation of mRNA metabolism” and “translation” still show up among the most significant terms, their significance is reduced (Fig 2D). Besides the SG associated proteins, we detected 51 significantly enriched proteins for MHV with no previous link to SG biology, 29 of which are also significantly enriched for the control condition (MHV APX+ vs. MHV APX-) (Fig 3B, S3). Besides cellular proteins, these factors also include 9 viral proteins (nsp1, nsp2, nsp3, nsp4, nsp8, 2a, M, S and N) (Fig 2B). Interestingly, our experiment revealed the significant depletion of 24 proteins from the G3BP1 microenvironment of MHV-infected cells. According to the GO term enrichment analysis, several of these proteins are associated with glycosylation (Fig 2D, middle). This posttranslational modification is known to be involved in cell signaling and was also previously linked to SG biology (26). As it was suggested that SGs might function as a signaling platform in the antiviral response (6,27), we checked for the presence of classical RNA sensors like RIG-I, MDA5 and PKR in infected/treated samples. However, none of these factors have been detected in the labelled SG proteome.

For the direct comparison between MHV and sodium arsenite induced stress, we observed enrichment of most SG associated proteins for arsenite-induced SGs similar to the mock comparison, although the fold change of most of the SG nucleator proteins dropped below our significance criteria (Fig 2C). Among the 115 significantly enriched proteins for arsenite, 27 have been identified as known SG associated factors (Fig 3C). In contrast, only 31 proteins are significantly enriched for MHV compared to arsenite, none of which were previously associated with SGs (Figs 2C/3C). Similar to the previous comparison, the MHV enriched SG proteome shows the presence of several viral proteins. Our analysis suggests that the arsenite induced stress response is stronger than the one induced by MHV. This is supported by the GO term enrichment analysis showing an enrichment for terms connected to mRNA metabolism, translation and SG formation for arsenite stress against MHV, however to a lesser extent than observed in the comparison with unstressed cells (Fig 2D). In contrast, the MHV infected condition, which also showed some enrichment for these terms in the previous mock comparison, is lacking their enrichment compared to arsenite stress.

Taken together, our proximity labelling screen showed the expected enrichment of SG associated proteins in the microenvironment of sodium arsenite induced SGs in line with previous findings. In contrast, the microenvironment of SGs during MHV infection revealed a decreased abundance of SG associated proteins and a general reduction in the protein variety. The two observed stress stimuli thus induce a distinct SG composition.

### SG are induced at distinct time points during MHV and SFV infection

The profound differences in stress granule proteomes between MHV infection and sodium arsenite treatment raised the question if the reduced abundance of several SG markers is a general feature of virus induced stress granules or rather an MHV- or coronavirus-specific occurrence. Thus, we extended our investigation to Semliki Forest virus (SFV), which was previously used in several studies to investigate stress granules in the context of virus infection and thus serving as a reference (9,15,28). First, we assessed the stress granule dynamics during MHV and SFV infection. For this purpose, we infected 17Cl1:G3AG cells with either MHV or SFV at a high multiplicity of infection (MOI=10) and repeatedly imaged the samples for up to 22hpi. For both infections, the formation of cytoplasmic accumulates of G3AG could be observed, although the timepoint of their appearance substantially differs between the viruses (Fig 4A, S1 movies). The quantification of the average SG number per cell displayed in figure 4B shows a rapid SG formation after 3hpi for SFV. The maximum SG number is reached between 4 to 5hpi, followed by a fast dissolvement of the condensates. At 8hpi almost all SGs disappeared, and no reformation can be observed for the rest of the experiment. This is in line with previous studies reporting the presence of SGs early in SFV infection and their subsequent dissolvement (9,15). For MHV infection, the SGs start to appear much later at 5hpi, reaching a peak at 8hpi as observed previously. Like for SFV, MHV-induced granules dissolve with progressing infection and almost no SGs can be seen after 14hpi. This delayed induction of SGs is in line with a previous study showing the presence of MHV-induced granules only after 6hpi (20).

**Fig 4.**
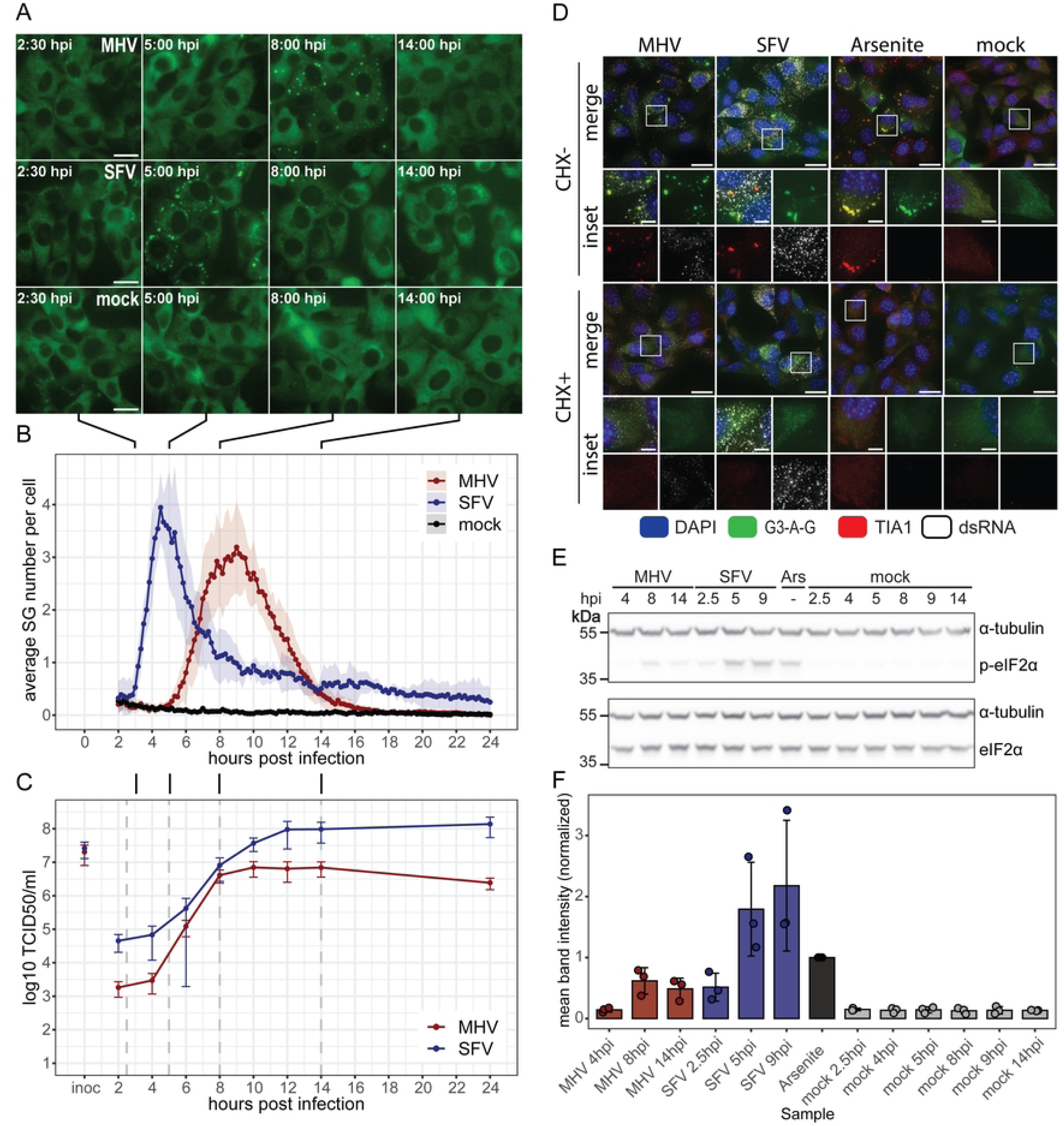
Analysis of SG formation during MHV and SFV infections by live-cell imaging. (A) 17Cl1 cells expressing a G3BP1-APEX2-GFP fusion protein (G3-A-G, green) were infected with either MHV or SFV at an MOI of 10, or mock infected. MHV infected samples were additionally supplemented with HR2 peptide to prevent syncytia formation. Cells were imaged every 10min for ∼22h on a DeltaVision widefield fluorescence microscope with on-stage incubator. The panel shows a selection of time points. (B) Quantification of average SG number per cell based on live-cell imaging data. SGs were segmented and counted using Ilastik and Fiji. Number of SGs per frame was standardized with the number of visible cells and the mean based on three separate image series for each condition was plotted in RStudio. The ribbon indicates the standard deviation. (C) Single cycle infection kinetics for MHV and SFV on 17Cl1:G3-A-G cells. Cells were infected with MHV or SFV at an MOI of 10. Supernatants were collected at the indicated time points and virus titers were estimated by TCID50 assay. (n = 3). (D) Effect of cycloheximide (CHX) on SGs during MHV and SFV infection or sodium arsenite treatment assessed by indirect immunofluorescence staining. 17Cl1:G3-A-G cells were either infected with MHV-A59 or SFV at an MOI of 10, or treated with sodium arsenite (250uM) to induce SGs. MHV infected samples were supplemented with HR2 peptide to prevent syncytia formation. CHX+ samples were treated with 50uM cycloheximide 1.5h prior to fixation. Cells were fixed at 8hpi (MHV), 5hpi (SFV) or 30min after sodium arsenite treatment. Cells were stained with antibodies against TIA1 (red) and dsRNA (grays) and imaged on a DeltaVision widefield fluorescence microscope. Scale bar: 20um or 5um (inset).

The SG induction timepoint might be dependent on the duration of the virus replication cycle. Thus, we next performed a one cycle replication kinetics for both viruses. 17Cl1 cells were infected at an MOI of 10 and the amount of newly formed infectious particles was measured at different time points. Both viruses have comparable replication kinetics showing an exponential release of new infectious particles after 4hpi before reaching a plateau within 12hpi (Fig 4C). Correlating the SG formation and replication kinetics, we find that the induction of SGs occurs early in SFV infection before the exponential growth and release of new virus. In contrast, MHV shows the SG formation at a later stage of infection simultaneously with its exponential growth phase. Interestingly, the granules persist during the whole exponential phase of MHV growth and are only dismantled once the viral titer reaches a plateau (Fig 4B/C). It was shown for several viruses that they form viral aggregates with SG components to interfere with the cellular stress response (8,29,30). To assess if MHV might form aggregates of SG markers disconnected from SGs, we performed cycloheximide treatment of infected cells. This drug stalls the ribosomes on the mRNA causing the dissolvement of canonical SGs, which are in an equilibrium with polysomes, a feature not observed for the viral aggregates (24,31). Without treatment we observe the formation of G3AG positive granules containing TIA1 for all three stress stimuli as seen previously (Fig 4D). In contrast, when cells are treated with cycloheximide no granules can be observed for the canonical stressor arsenite, nor for any of the viruses (Fig 4D). Based on these results, MHV- and SFV-induced granules are in equilibrium with polysomes like canonical SGs contradicting the viral accumulations in MHV infection.

The major pathway of canonical SG induction is the integrated stress response (ISR). For SFV it is well known that ISR is activated early in infection leading to SG formation (15). To investigate, if SG formation during MHV infection occurs together with ISR activation, we next analyzed the level of phosphorylated eIF2α (p-eIF2α), the main marker for ISR activation, at different time points during MHV and SFV infection. We further used sodium arsenite treatment as a positive control for canonical stress. The Western blot analysis reveals greatly increased levels of p-eIF2α for all SFV time points, whereby the later time points exceed the level of arsenite stress (Fig 4E/F). A small increase in p-eIF2α level can also be seen for MHV at later time points indicating the activation of ISR (Fig 4E/F). However, it is substantially lower than our positive control.

Collectively, our data showed the formation of SG-like condensates during MHV and SFV infections, which are in equilibrium with the translation machinery similar to canonical SGs. Simultaneously with SG appearance, both viruses induce the activation of the ISR, although MHV shows a much weaker activation compared to SFV and arsenite. Furthermore, the induction timepoint of these condensates in relation to the replication cycle differs strongly between the viruses.

### G3BP1 microenvironment diverges between MHV and SFV infected cells during SG formation

Next, we comprehensively compared the SG proteome between MHV and SFV infection to assess if previously observed characteristics of MHV-induced SGs are also observed for other viruses. We again applied APEX2-mediated proximity labelling in combination with SILAC. To control for virus-induced changes in G3BP1 microenvironment in the absence of SGs, the labelling reaction was conducted at three different time points in relation to SG formation. Thereby time points before SG induction (pre SG, Fig 5A), at SG peak (peak SG, Fig 5B) and after SG dissolvement (post SG, Fig 5C) were selected for MHV and SFV and directly compared between both viruses. The proximity labelling was verified by immunofluorescence and Western blot analysis (S4 Fig A/B). Furthermore, the labelling control condition showed a strong enrichment of most proteins for the unstressed proximity labelled condition compared to the unlabeled condition in line with a successful proximity labelling (S5 A). We detected 3502 different proteins over all analyzed samples, 230 of which are SG associated proteins based on our reference list. Hereafter we focus primarily on proteins detected in both compared conditions, thus yielding a heavy/light ratio.

**Fig 5.**
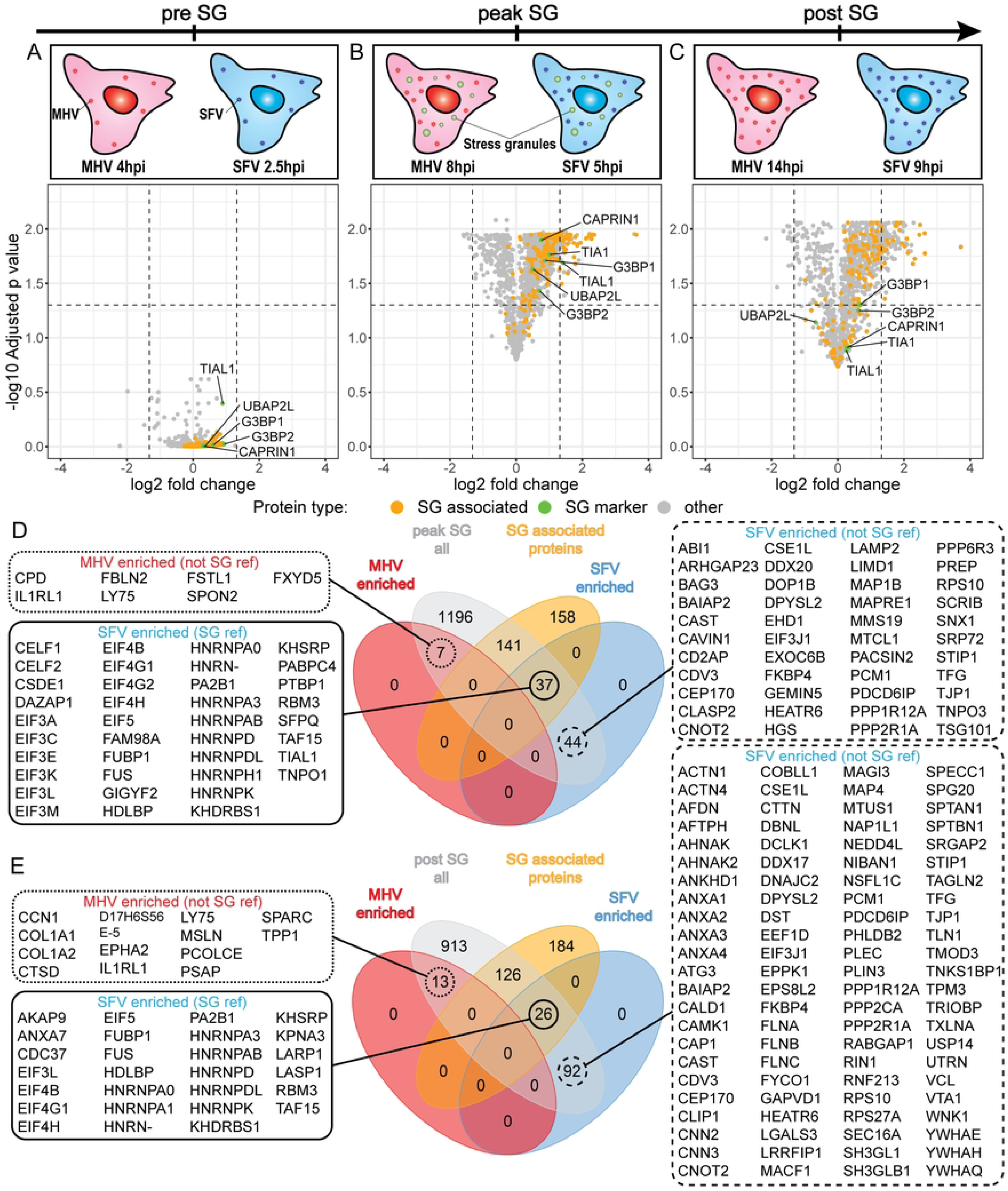
Direct comparison of the G3-A-G microenvironment between MHV and SFV infection at different time points. Scheme visualizing the timeline and the compared conditions. (A) Different abundance of proteins between MHV-infected (4hpi) and SFV-infected (2.5hpi) and proximity labelled cells prior to SG formation. (B) Different abundance of proteins between MHV-infected (8hpi) and SFV-infected (5hpi), proximity labelled cells at the respective peak SG time point. (C) Different abundance of proteins between MHV-infected (14hpi) and SFV-infected (9hpi), proximity labelled cells after SG disappearance. Volcano plots display the mean log2 fold change based on the heavy/light protein ratio between the conditions (n = 3) plotted against the -log10 adjusted p value. Proteins contained in the reference list for SG associated proteins are indicated in orange, the major SG markers are labeled and indicated in green. Dashed lines show the significance criteria for log2 fold change >log2(2.5) and <-log2(2.5) respectively, and adjusted p value < 0.05. Venn diagrams visualizing overlap between significantly enriched protein fractions (red: MHV / blue: SFV), all identified proteins (gray) and reference list of SG associated proteins (orange) for (D) the peak SG time point and (E) the time point after SG disappearance (post SG). For selected sections the corresponding lists of proteins are provided.

For the comparison of the pre SG time point between MHV and SFV we detected 1990 proteins in both conditions, whereby 186 were previously associated with SGs including several major SG markers (Fig 5A, S5 C). Interestingly, no significant differences in protein abundance can be observed between the viruses based on our significance criteria (log2(2.5) and p 0.05). These results indicate that the G3BP1 microenvironment seems to be similar in both virus infections prior to SG formation.

In contrast to the pre SG timepoint, we detected substantial changes in the G3BP1 microenvironment between MHV and SFV at the peak of SG presence. We found 81 proteins to be significantly enriched for SFV-induced granules compared to MHV (Fig 5B). Almost half of these proteins (37) belong to our list of SG associated proteins indicating a strong connection to SG biology of the SFV-enriched fraction (Fig 5D). Furthermore, we found a tendency of most SG associated proteins, including the major SG markers, to be enriched for SFV compared to MHV as seen in the volcano plot and the log2 fold change density distribution of SG associated proteins (Fig 5B, S5 B). In contrast, only 7 proteins are significantly enriched for MHV-induced granules compared to SFV, none of which was previously associated with canonical SGs (Fig 5B/D). Based on these results, SFV-induced granules match very well to canonical SG components while the MHV-induced ones show substantial differences. These findings are in line with an additional SILAC experiment directly comparing SFV-, MHV- and arsenite-induced SGs with unstressed mock cells. Similar to arsenite-induced SGs, the granules formed during SFV infection show an enrichment for known SG associated proteins including the major markers G3BP1/2 (S6 Fig A/C, S7 Fig A/C). For MHV we can observe a high log2(FC) enrichment for the major SG nucleator (S6 Fig B, S7 Fig B). But the majority of SG associated proteins show a lower fold change. Furthermore, the analysis shows a higher uncertainty as none of the proteins surpasses the p value criteria <0.05. These results indicate a less profound change of the G3BP1 microenvironment compared to the unstressed state.

After SG dissolvement (post SG time point), the microenvironment of G3BP1 is still characterized by differences in protein composition between MHV and SFV infection (Fig 5C). However, despite an increased number of significantly enriched proteins for SFV compared to the peak SG time point, the number of SG associated proteins (26/118) decreased in line with the dissolvement of SGs (Fig 5C/E). As many of these factors are not overlapping with the enriched fractions at the peak time point, the differences in G3BP1 microenvironment might represent general virus-specific changes in the cellular state or the accumulation of G3BP1 in other structures. For both viruses we detected viral proteins foremost in the peak and post SG conditions (S8 Fig). MHV showed the same set of proteins detected in the previous SILAC experiment. For SFV, all non-structural proteins have been detected with nsP3 as the most highly enriched one, which is in line with its G3BP1 interaction (9,29).

Based on the Venn diagrams, we further investigated subsets of significantly enriched proteins for the peak and post time points provided as protein lists in figure 5D/E. Among the significantly enriched SG associated proteins for SFV peak SG we identified two dominant groups of proteins, the first consisting of several translation initiation factor (eIF) subunits and the second consisting of different members of the heterogeneous nuclear ribonucleoproteins (hnRNP) family (Fig 5D). Interestingly, many of the hnRNP members are amongst the most strongly enriched hits for SFV with a fold change up to 8 for FUS (Fig 5B).

To functionally classify the sets of significantly enriched proteins in the peak and post comparisons, we performed a GO term enrichment analysis. The enriched protein fraction for SFV at the SG peak time point revealed “regulation of mRNA metabolic process” and “translation initiation” as most significant terms for biological process (Fig 6A). A closer look at the associated proteins for both terms showed the two previously mentioned groups of hnRNPs and eIFs as predominant proteins (Fig 6D/E). Consistently, the analysis for cellular compartment (CC, Fig. 6B) and molecular function (MF, Fig 6C) revealed a strong link to ribonucleoprotein granules, translation initiation factor 3 complex, translation regulation activity and mRNA binding. Most of these GO terms are also found for the SFV post SG enriched proteins (Fig 6A/B/C). However, their associated count is reduced, which is in line with the reduced number of eIFs and hnRNPs enriched for the SFV post SG condition (Fig 5E). For MHV we only found a few significantly enriched proteins for peak and post time points strongly limiting the meaningfulness of the GO term enrichment analysis for these samples (Fig 6A/B/C).

**Fig 6.**
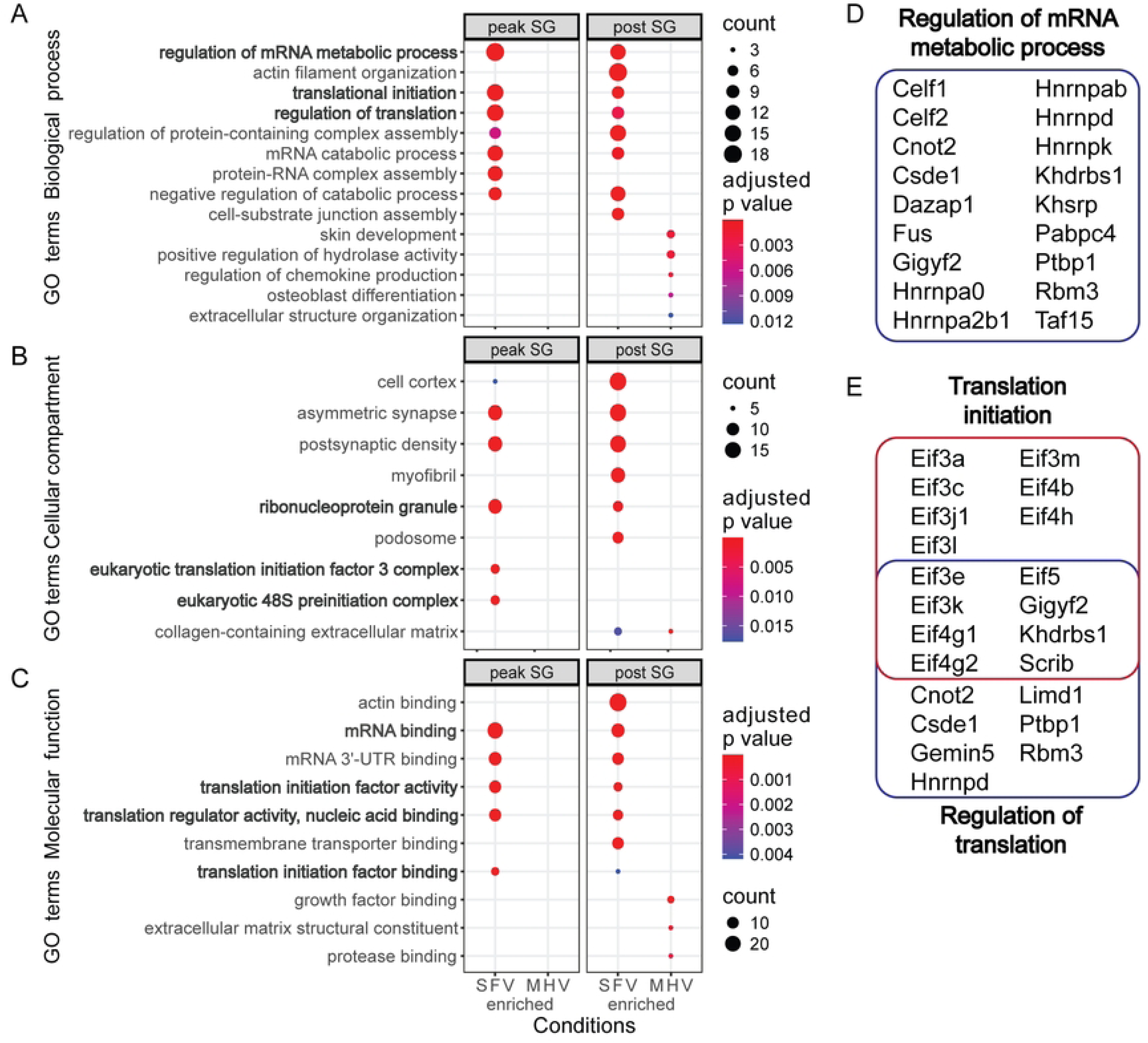
Functional analysis of significantly enriched/depleted proteins for SFV- vs. MHV-induced SGs. Subsets of significantly enriched proteins for SFV and MHV at the peak and post SG time points based on the direct comparison (|log2FC| > log2(2.5), adjusted p value < 0.05) were subjected to a GO term enrichment analysis using ClusterProfiler. Redundant terms for each condition were reduced based on semantic similarity. Displayed are the top 5 most significant terms found in each subset of proteins for (A) biological processes, (B) molecular function and (C) cellular compartment. The circle size indicates the number of identified proteins for each term. Color indicates adjusted p value based on Benjamini–Hochberg. Terms of special interest in the context of SGs are highlighted in bold. Lists of proteins associated with the biological process (D) regulation of mRNA metabolic process, and (E) translation initiation, regulation of translation identified in this screen.

These results provide a detailed insight into the G3BP1 microenvironment at different time points during MHV and SFV infections by directly comparing both viruses. Prior to SG formation the G3BP1 microenvironment is similar for both viruses. When SGs are present, we observe substantial virus-specific differences characterized by an enrichment of many SG associated proteins including several members of eIFs and hnRNPs for SFV-induced SGs compared to MHV. Virus-specific differences in G3BP1 microenvironment also persist after dissolvement of SGs. Overall, in the direct proteome comparison between MHV- and SFV-induced SGs as well as the comparisons with mock, SFV reveals an increased connection to canonical SG biology.

### Reduced abundance of specific SG markers in MHV-induced SGs compared to SFV

Next, we aimed to verify selected proteins identified as significantly enriched for SFV-induced SGs by investigating their subcellular localization in SFV and MHV infected cells at the SG peak time point. Therefore, we applied indirect immunofluorescence microscopy on 17Cl1:G3AG cells staining for several eIFs (eIF3a, eIF3e, eIF4G1, eIF4H) and proteins involved in mRNA processing (FUS, PTBP1, TAF15). Additionally, we also stained double-stranded (ds) RNA as an intermediate of viral replication to visualize the presence of the viruses. Besides MHV and SFV infections, sodium arsenite treatment was included as reference for canonical SGs. All investigated translation initiation factors showed a strong accumulation in G3AG positive granules for the sodium arsenite treated samples, while they remained diffusely distributed in our mock control (Fig 7A-D, bottom row). A similar accumulation of eIFs and colocalization with G3AG positive granules can be observed in cells infected with SFV (Fig 7A-D, middle row). In contrast, the probing for eIFs in MHV infected cells only shows a weak accumulation of these proteins in G3AG positive granules (Fig 7A-D, top row). Thus, the previously observed difference in the abundance of eIFs between MHV- and SFV-induced SGs suggests a mechanism for exclusion or reduction of these proteins from MHV-induced granules, while the SFV-induced SGs are similar to canonical SGs induced by arsenite. The staining for the major SG markers, G3BP1, TIA1 and UBAP2L, as well as eIF4A3 revealed a similar accumulation for all investigated SGs (S9 Fig).

**Fig 7.**
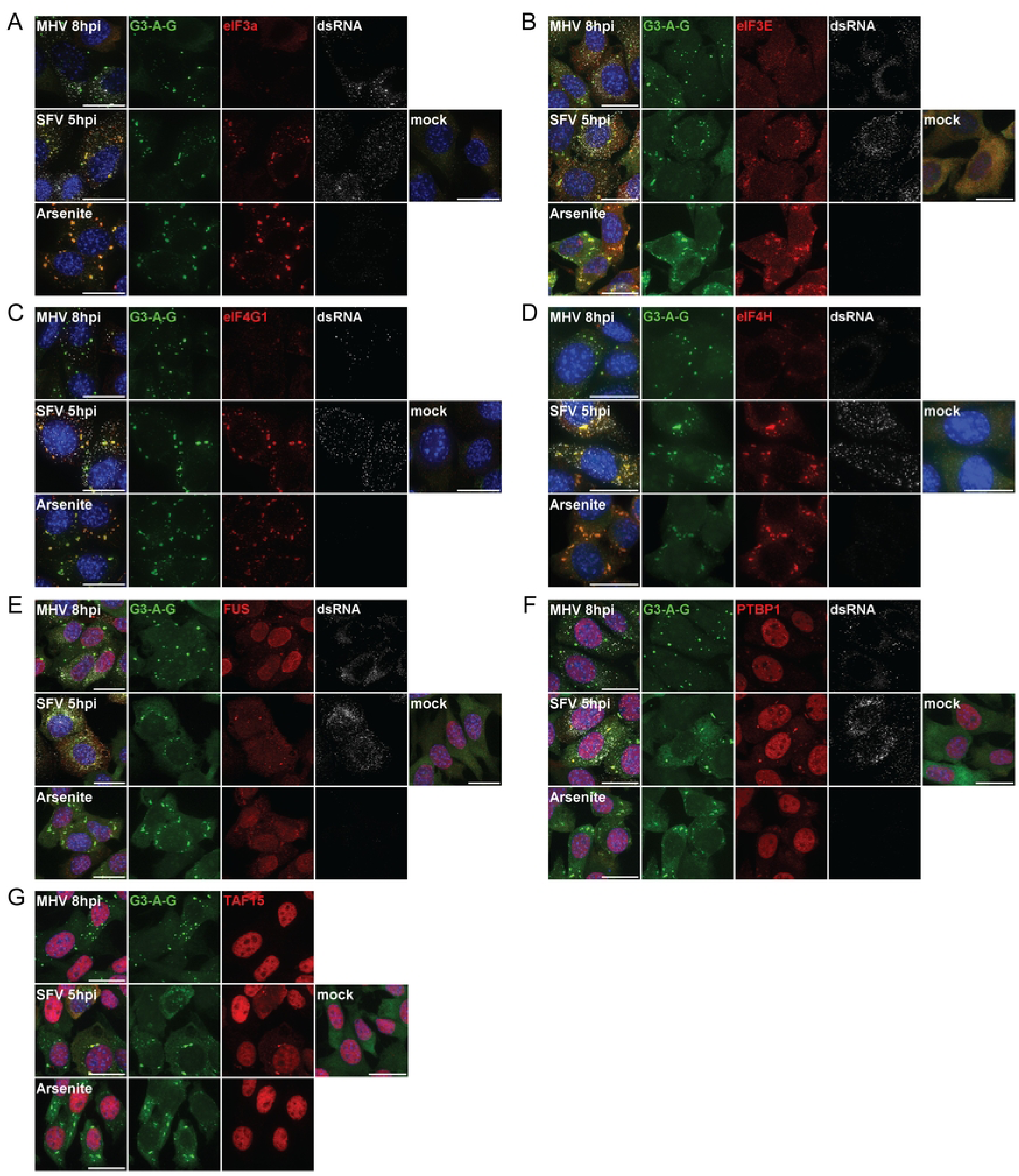
Subcellular localization of SG proteins during different virus infections or oxidative stress by indirect immunofluorescence. 17Cl1:G3-A-G cells (green) were infected with either MHV or SFV at an MOI of 10, mock infected or treated with 250uM sodium arsenite for 30min. Cells were fixed at the respective peak time point of SGs and subjected to indirect immunofluorescence staining. Staining of nucleus with DAPI (blue). (A) Staining of eIF3a (red) and dsRNA (grays). (B) Staining of eIF3L (red) and dsRNA (grays). (C) staining of eIF4G1 (red) and dsRNA (grays). (D) Staining of eIF4H (red) and dsRNA (grays). (E) Staining of FUS (red) and dsRNA (grays). (F) Staining of PTBP1 (red) and dsRNA (grays). (G) Staining of TAF15 (red). All MHV-infected samples were supplemented with HR2 peptide to prevent syncytia formation. Images were acquired on a DeltaVision widefield system or a Nikon spinning disk confocal system and processed in Fiji. Scale bar: 20um.

The staining for specific hnRNP members revealed a different picture. While a weak accumulation of FUS, PTBP1 and TAF15 can be observed in arsenite and MHV induced G3AG granules, the majority of these proteins still localizes to the nucleus similar to the mock control (Fig 7E-G). For SFV infected cells we found two different phenotypes. Some cells show a predominant localization of FUS, PTBP1 and TAF15 in the nucleus with only a weak accumulation in SGs if present. Some cells, however, display an even distribution of these proteins throughout the whole cell, which comes together with a stronger accumulation in virus-induced SGs (Fig 7E-G). Many cells with delocalization of FUS to the cytoplasm also display an intense signal for dsRNA, indicating a progressed stage of infection. To further verify this observation, we performed a time course experiment staining SFV-infected cells at different time points after infection. The number of cells with cytoplasmic localization of nuclear proteins increases later in infection together with an increased dsRNA signal (S10-12 Figs).

When considered together, the subcellular localization of various proteins by indirect immunofluorescence microscopy verified the virus-specific differences in SG protein composition. We confirmed the difference in abundance of several eIFs in MHV-induced granules while the SFV-induced ones are similar to canonical SGs. In contrast, the enrichment of several hnRNPs seems to be specific for SFV-induced SGs and is likely connected to delocalization of these proteins with progressing infection. This observation highlights another virus-specific characteristic of the investigated granules.

### MHV genomic and subgenomic RNA is not accumulated in SGs

Besides proteins, RNA is a major component of SGs and it was previously observed that not only cellular mRNA but also viral mRNA can accumulate in these condensates (5,32). Thus, we wondered if viral mRNA is accumulated in SGs induced by MHV or SFV. To assess the subcellular distribution of viral RNA, we applied single molecule RNA fluorescence in situ hybridization (smRNA-FISH). This technique relies on a pool of fluorescently labelled probes complementary to the sequence of interest allowing single molecule resolution (33). To visualize MHV mRNA we designed two different probes. One targets the nsp3-coding sequence in the ORF1a and thus only detects full length viral genomes (Fig 8A). The second probe binds the sequence of the spike protein staining full length genomic RNA and subgenomic mRNA of the spike gene. Both probes in combination allow to distinguish between genomic and subgenomic RNA of MHV (Fig 8A). For SFV we designed one probe targeting the ORF1 (nsP2-coding region) to detect full length genomic RNA (Fig 8A).

**Fig 8.**
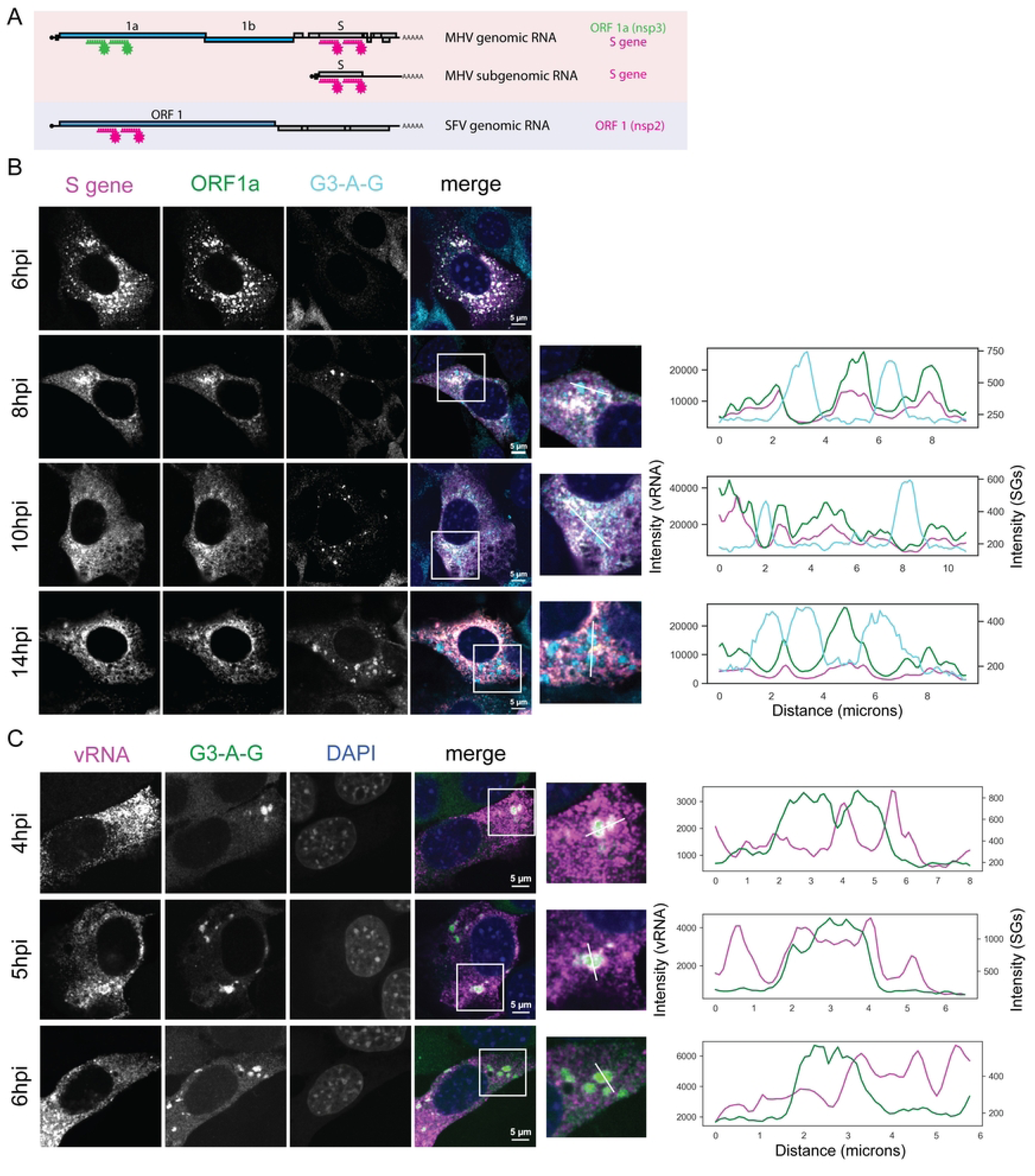
Subcellular localization of viral RNA in relation to SGs during MHV and SFV infection using smRNA-FISH. (A) Scheme indicating the different probes and their binding location within the MHV or SFV genome or subgenomic RNA. To visualize genomic and subgenomic RNA of MHV two different probes were designed, each consisting of a pool of 50 different, fluorophore-conjugated oligonucleotides complementary to either the nsp3 gene or the S gene. For staining of SFV genomic RNA a probe targeting the nsp2 gene was designed. (B) Visualization of MHV genomic (double positive) and subgenomic (positive for S probe) RNA in relation to SGs at different time points after infection. 17Cl1:G3-A-G cells (cyan) were infected with MHV-A59 at an MOI of 0.1. After infection, medium was supplemented with HR2 peptide to prevent syncytia formation. Cells were fixed at the indicated time points and stained for viral RNA using probes targeting the nsp3 gene (green) and the S gene (magenta). (C) Visualization of SFV genomic RNA in relation to SGs at different time points after infection. 17Cl1:G3-A-G cells (green) were infected with SFV (MOI 0.1) and fixed at indicated time points. FISH staining was performed using probe (magenta) complementary to SFV nsP2 gene. All images were acquired with a Leica spinning disk confocal microscope and post-processed in Fiji. Intensity line plots display the measured signal intensity of the different probes and the G3-A-G along the white line indicated in the inset. Scale bar: 5um

17Cl1:G3AG cells were infected with either MHV or SFV and different time points around the previously used SG peak time point were selected for both viruses to be analyzed. The resulting images for MHV can be seen in figure 8B. An intense signal for both vRNA probes can be observed at all time points. At 6hpi well defined foci in the perinuclear region can be observed, which are positive for both probes, indicating genomic viral RNA. These foci likely represent the replication organelles. In all later time points, the staining for vRNA becomes more evenly distributed throughout the cytoplasm (Fig 8B). Due to the high amount of vRNA present at all investigated time points no individual RNA molecules can be resolved. G3-A-G positive cytoplasmic foci are present in infected cells of all but the 6hpi time point. Interestingly, the G3AG fusion protein shows an inverse correlation with both viral RNA probes for all cells with SGs. This is evident in the intensity line plots (Fig 8B), revealing absence of vRNA at sites of high G3AG signal representing SGs. A staining for an endogenous mRNA, the activating transcription factor 4 (ATF4) previously shown to accumulate in SGs (34), revealed that in contrast to the viral RNA cellular mRNA is readily detectable in MHV-induced SGs (S13 Fig). These findings indicate that MHV genomic and subgenomic mRNAs are excluded from virus-induced SGs.

Similar to MHV, cells infected with SFV show the presence of large amounts of viral genomic RNA in the cytoplasm for all investigated time points (Fig 8C). In contrast to MHV, genomic RNA of SFV is not excluded from virus-induced SGs at all imaged time points. Even more, the viral RNA seems to have accumulated in the G3AG positive foci. As visible in the intensity line plots, this accumulation concentrates particularly at the edge of the SGs (Fig 8C). Collectively, the subcellular localization of viral RNA in relation to SGs revealed virus-specific differences in SG characteristics between MHV and SFV infection also on the RNA level. While for SFV the genomic viral RNA is accumulated mainly at the edges of virus-induced SGs, MHV revealed an efficient exclusion of its genomic and subgenomic mRNAs from these condensates. Overall, our investigation into various aspects of SG formation during MHV and SFV infection showed that these condensates are largely characterized by the type of virus inducing them.

## Discussion

A major focus of this study was based on characterizing the protein composition of the virus-induced SGS, as the presence or absence of specific proteins provides insight into possible roles of these condensates and associated host-virus interactions. For oxidative stress-induced SGs, which we use as a positive control, our results showed the expected proteome of canonical SGs. In line with previous findings by Markmiller et al. and others, we identified the major SG nucleator proteins, components of the preinitiation complex and a large variety of other proteins involved in RNA metabolism as abundant components of oxidative stress induced SGs verifying the labelling of the SG microenvironment (12,13). Our screen further provides a set of novel SG protein candidates. Interestingly, we also detected several known SG associated proteins in the G3BP1 microenvironment under unstressed condition as seen in the proximity labelling assay. A similar observation was made by Markmiller et al. and others suggesting that this could indicate a preexisting network of SG protein interactions in the unstressed cell, which is intensified upon stress (12,13).

The granules formed during MHV infection show the accumulation of the major SG nucleator proteins including G3BP1, TIA1 and UBAP2L similar to canonical SGs. The presence of these SG markers was similarly observed in previous studies (20). Several viruses, like Japanese encephalitis virus, have been shown to accumulate SG proteins in distinct viral aggregates to prevent SG formation (8). However, this does not seem to be the case for MHV induced granules as they are in an equilibrium with polysomes indicated by their dissolvement with polysome stabilizing compounds. Furthermore, the increase in phosphorylated eIF2α simultaneously with granule appearance suggests that they are a result of the activated stress response during late MHV infection. Despite these canonical SG properties, MHV-induced SGs show a lower abundance of the majority of SG associated proteins including eIFs, and an overall reduced connection to SG biology compared to arsenite induced SGs. Thus it is tempting to speculate that MHV induces a weaker stress response with less profound changes to the G3BP1 microenvironment and may even employ mechanisms to exclude or greatly reduce abundance of many canonical SG-associated proteins. In line with this, MHV infected cells show a lower level of phospho-eIF2α compared to arsenite stress.

In accordance with previous studies, we found the SGs formed during SFV infection resemble canonical SGs. They are formed upon the induction of the ISR by PKR activation and have a high abundance of SG nucleator proteins and components of the translation initiation machinery (15,28). In our study, the ISR activation even exceeds the level of arsenite induced stress, demonstrating an efficient mounting of the stress response by infected cells early in SFV infection. Comparing the SG microenvironment between SFV and MHV, we found a similar enrichment of SG associated proteins for SFV as seen for oxidative stress, suggesting that the observed depletion of these factors for MHV induced SGs is a virus-specific characteristic. Thus overall, SFV-induced SGs seem to be more similar to canonical SGs than MHV-induced ones. The most striking difference thereby is a strong depletion of several translation initiation factors from SGs in MHV-infected cells, which seemingly exceed the reduced abundance of the other SG markers. Translation initiation factors are abundant in canonical SGs as most of them are part of the 48S preinitiation complex, a major component of SGs (35). Furthermore, many of these proteins are important host factors for capped RNA viruses like coronaviruses and alphaviruses, as they rely on the host translation machinery for viral protein production. It was suggested that SGs might exert antiviral properties by sequestering eIFs to inhibit viral translation (36). Based on our results, the eIFs seem to be either actively or passively prevented from accumulating in SGs during MHV infection keeping them available for viral translation. The different eIF abundance in MHV and SFV induced SGs thus might indicate different ways how the viruses cope with the stress reaction and SG formation and ultimately a different role of these granules in both virus infections.

Besides the distinct protein composition our study further revealed virus-specific differences in the accumulation of vRNA in MHV- and SFV-induced SGs. This is of special interest because some suggested antiviral mechanisms of SGs are associated with the recruitment of vRNA to these condensates. By bringing together vRNA and sensors like RIG-I and PKR, SGs could serve as signaling platforms regulating the antiviral response (5,7). However, in contrast to several studies reporting an accumulation of RIG-I, PKR and OAS in SGs during Sendai virus and influenza A ΔNS1 virus infections, our proteomics screen showed no specific enrichment of these proteins neither for MHV, nor for SFV (5–7). Another suggested antiviral mechanism is the sequestration of vRNA as a recent study found that the condensation of vRNA in SGs during SARS-CoV-2 infection limits viral growth (10). In the context of these findings, it is striking that MHV efficiently excludes gRNA and sgRNA from virus-induced SGs, while SFV shows a strong accumulation of gRNA in these granules. Because the observed exclusion of vRNA for MHV could counteract the proposed antiviral mechanisms of SGs, our findings suggest different roles of the granules during MHV and SFV infections. However, if this exclusion is based on an active mechanism or rather on the efficient sequestration of MHV RNA in the replication organelles must be further elucidated.

Besides the differences in composition, the varying role of MHV and SFV induced SGs during both infections is further supported by their formation kinetics observed in our live-cell imaging. SFV triggers the SG formation early in infection similarly as observed previously (9). This formation is preceded by a strong activation of the cellular stress response seen by elevated phospho-eIF2α levels already before SG assembly. With progressing infection SFV actively disassembles the granules likely mediated by the nsP3 directly interacting with G3BP1 (9,32). This dissolvement of SGs seems to occur prior to the exponential growth phase of SFV suggesting that SFV might have to disassemble SGs to facilitate production of virus progeny, which is in line with a previous study reporting a correlation between SG disassembly and higher levels of viral RNA (28). In contrast, the SGs during MHV infection only appear at a late stage of infection with increasing production of new virus particles. At early stages of infection, the virus seems to prevent the induction of the stress response and subsequent SG formation. If this inhibition is based on an active mechanism or rather an efficient hiding of MHV from cellular sensors must be further investigated. However, the latter is likely, as it is well known that coronaviruses hide their vRNA in double membraned vesicles and further possess an arsenal of nsps, like nsp15, to prevent its detection by host sensors (18). This might also explain the absence of vRNA in MHV-induced SGs observed in our study. When the granules are finally formed the virus has already established the infection and completed most of its live cycle. Thus, MHV does not only seem to induce a weaker stress response compared to SFV, but it also does so only at a late stage where a potential antiviral effect on viral growth is limited.

Altogether, our study revealed substantial virus-specific differences for the investigated SGs contradicting the formation of a general “antiviral SG”. It further supports different roles of these granules during SFV and MHV infection strongly shaped by different virus-host interactions. Our current understanding (Fig 9) is that SFV induces the formation of canonical SGs early in infection likely triggered by sensing of vRNA via the ISR. These SGs accumulate canonical SG proteins including eIFs as well as viral RNA, which could limit viral growth by reducing the availability of these components for SFV. To efficiently produce new virus particles, SFV actively dissolves the granules as infection progresses and prevents their reformation (9). In contrast, MHV prevents ISR activation and SG formation during the early phase of infection by efficiently hiding its vRNA. Only at a later stage is the stress response activated and G3BP1 positive granules are formed. However, these granules show a reduced abundance of many SG proteins and an altered composition distinct from canonical SGs. The reduced accumulation of eIFs and the exclusion of vRNA from MHV-induced granules likely counteracts antiviral functions mediated by sequestering important host-factors and vRNA. The altered composition in combination with the late induction time point allows the virus to tolerate the presence of these granules until their dissolvement during the collapse of cellular homeostasis. Collectively, our work revealed a remarkable and surprising plasticity of SG formation and composition that is to a large extent dependent on the type of virus infection. In particular the greatly reduced abundance of canonical SG proteins in MHV-induced SGs and the exclusion of MHV RNA from SGs raise the question, if these granules exert any antiviral function. For example, if MHV, and possibly other coronaviruses, employ mechanisms (i) to prevent SG formation early during infection, (ii) to modulate the SG proteome, and (iii) to exclude viral RNA from SGs. Our work also provides an impetus for future studies on SG biology in general, and the role(s) of SGs during infection with coronaviruses, alphaviruses and beyond. As with many other cellular processes, basic principles of cell biology can be deciphered by using virus infection models, and as our study suggests, basic determinants of SG formation and SG composition are still unknown.

**Fig 9.**
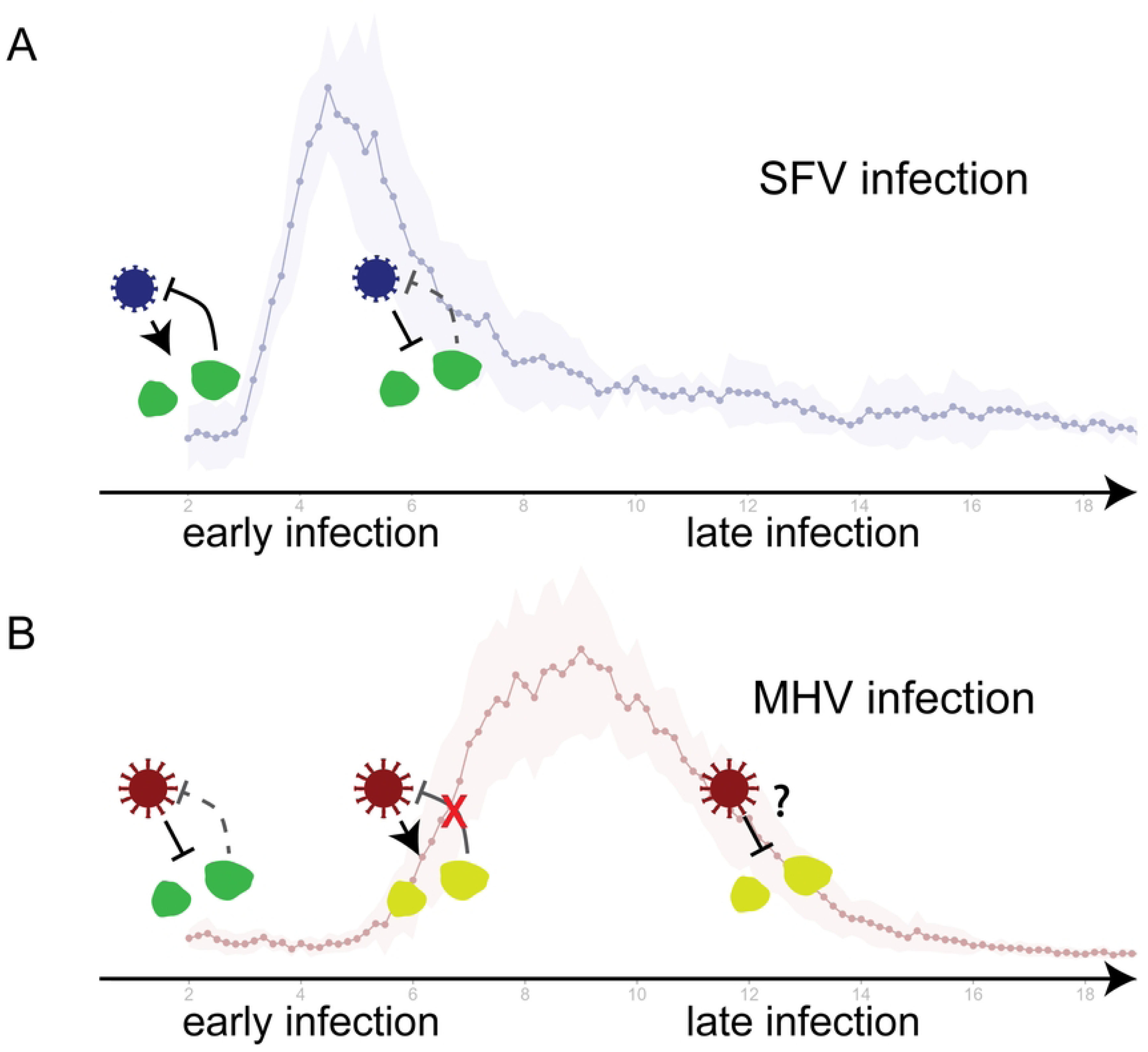
Proposed model of host-virus interaction in the context of SGs during SFV and MHV infections in cells. SFV seems to induce canonical SGs early in infection characterized by the presence of well-known SG marker proteins like eukaryotic translation initiation factors. The sequestration of especially the eIFs, which are required by the virus for efficient viral translation, as well as the accumulation of SFV genomic RNA in these condensates could have negative effects on viral growth. Thus, the virus dissolves the SGs with progressing infection. In contrast, MHV seems to prevent induction of canonical SGs early in infection. Only with the onset of exponential release of new virus particles, accumulations positive for the major SG markers G3BP1, TIA1 and UBAP2L are formed in the cytoplasm. A reduced abundance of many known SG proteins including eIFs compared to canonical SGs suggests a type of atypical SGs. Together with the efficient exclusion of MHV viral RNA, this gives rise to the apparent tolerance of these condensates by MHV later in infection.

## Material and Methods

### Cell culture

Murine 17 Clone 1 fibroblasts (17Cl1, kindly provided by S.G. Sawicki) and murine 929 fibroblasts (L929, Sigma) were cultured on minimum essential medium (MEM, Gibco) supplemented with 10% (v/v) FBS, 1x Penicillin-Streptomycin (seraglob) and 1% (w/v) non-essential amino acids (Gibco)(MEM+/+/+). Human Hek293T cells were cultured in Dulbecco’s Modified Eagle Medium-GlutaMAX (DMEM, Gibco) supplemented with 1 mM sodium pyruvate, 10% (v/v) FBS, 100μg/ml streptomycin, 100 IU/ml penicillin (Gibco) (DMEM+/+). Cells were grown in a humidified incubator at 37°C and 5% CO2.

### Viruses

Recombinant mouse hepatitis virus (MHV) A59 strain was previously generated using a reverse genetic system based on vaccinia virus(37). A plasmid encoding the genome of Semliki Forest virus (SFV) was kindly provided by Oliver Mühlemann. The sequence was modified with standard molecular techniques to represent the wildtype SFV strain 4. SFV mRNA was in vitro transcribed from the linearized plasmid and capped using the RiboMAX^TM^ Large Scale RNA Production System for the SP6 promoter (Promega). To rescue infectious virus, baby hamster kidney cells (BHK21) were electroporated with SFV mRNA and cocultured with susceptible 17Cl1 cells. The resulting passage 0 was used to produce progeny passage stocks on 17Cl1 cells for all infection experiments. Viral titers were estimated by plaque assay on 17Cl1 or L929 cells. To summarize, cells were grown in 24-well plates and infected with a 10-fold serial dilution of virus stock. The infection medium was replaced with overlay medium (DMEM supplemented with 10% FBS, 1x P/S, 1x NEAA and 1% methylcellulose) 2hpi. After one day, the cells were fixed with 4% formalin for 15min and stained with crystal violet for 10min. Titers were calculated based on the counted plaques.

### Generation of stable cell lines 17Clone1: G3BP1-APEX2-eGFP cells

First, the In-Fusion HD cloning kit (Takara) was used to integrate the coding sequence of the fusion protein mG3BP1-APEX2-GFP under the control of an HSV-TK promotor into an adapted pSMWUP lentiviral vector (CellBioLabs). Next, lentiviruses were produced in Hek293T cells by co-transfecting the cells with modified pSMWUP(TK:mG3BP1-APEX2-eGFP), pMD2G (adgene 12259) and CMVR8.74 (addgene 22036) using X-tremeGene HP DNA Transfection Reagent (Sigma-Aldrich). Produced lentiviruses were used to infect 17Cl1 cells, which were subsequently treated with puromycine (2ug/ml) to select for transduced cells. Surviving cells were sorted based on GFP signal using flow cytometry. 17Cl1:G3BP1-APEX2-eGFP (17Cl1:G3-A-G) cells were further propagated in MEM+/+/+ maintaining the puromycin selection at 1ug/ml. Puromycin was omitted for all experiments.

### Viral replication kinetics

17Cl1 cells were seeded in 24-well plates to reach 80-100% confluency the following day. Cells were infected with MHV or SFV at an MOI of 10 or mock infected without virus. Infection was synchronized by incubation of cells on ice for 1h, followed by incubation at 37°C for 1.5h. Cells were washed 3x with PBS and fresh MEM+/+/+ was added. Supernatant was collected at time points indicated in figure 4C and virus titers were estimated by TCID50 assay. The kinetics time course was carried out in three independent replicates. The mean titers and corresponding standard deviations were calculated and plotted in RStudio using ggplot2 and dplyr packages (38).

### Indirect immunofluorescence

Glass coverslips were coated with poly-D-lysine solution for 1h and washed 3x with PBS. Recombinant 17Cl1:G3-A-G cells were seeded on coated glass coverslips in a 12-well plate at a density of 2.25x10^5^ cells/well. The following day, cells were infected with MHV or SFV at an MOI of 10 and infection was synchronized as described above. Cells were washed 3x with PBS and fresh MEM+/+/N was added. Medium for MHV samples was supplemented with 1uM HR2 peptide. Samples for sodium arsenite treatment were incubated with MEM+/+/N supplemented with 250uM NaAsO_2_ (Sigma) 30 min before fixation. For cycloheximide treatments, the medium was exchanged 1.5h before fixation with medium supplemented with 50uM cycloheximide (Sigma). At the indicated time point, cells were washed 2 times with PBS and fixed with 4% formalin for 20min. Next, cells were washed and stored in PBS at 4° until staining.

For the staining procedure, the cells were permeabilized first by incubation in 0.1% Triton X-100 (in PBS) for 5 min at room temperature, followed by 20 min incubation in confocal buffer (CB) consisting of PBS supplemented with 50mM NH4Cl, 0.1% saponin, and 2% BSA. Afterwards, primary antibodies were diluted in CB as indicated in table 1. Glass coverslips were incubated in primary antibody solution for 2h at room temperature. Excess antibody solution was removed by washing the cells 3x 5min with CB, before incubating the cells with secondary antibody solution for 1h at room temperature. Afterwards, samples were washed firstly 3x with CB for 5min, followed by one wash with PBS and a quick rinse with water. Lastly, the cells were mounted on ProLong® Diamond mounting medium containing DAPI (Lifetechnologies).

**Table 1:** Antibodies/reagents for indirect immunofluorescence staining and Western blot analysis with used dilution factor and source.

| Antibody or reagent | Source | Identifier | Used dilution |
| --- | --- | --- | --- |
| anti-dsRNA (mouse mAb) | SCICONS | J2 | 1:200 |
| anti-G3BP1 (rabbit pAb) | Sigma | G6046 | 1:200 |
| anti-G3BP1 (mouse mAb) | Proteintech | 66486-1-Ig | 1:200 |
| anti-TIA1 (rabbit pAb) | Proteintech | 12133-2-AP | 1:200 |
| anti-UBAP2L (rabbit mAb) | Cell signalling | 40199 | 1:200 |
| anti-eIF3a (rabbit mAb) | Cell signalling | 3411 | 1:400 |
| anti-eIF3E (rabbit pAb) | Proteintech | 10899-1-AP | 1:50 |
| anti-eIF3L (mouse mAb) | proteintech | 67719-1-Ig | 1:400 |
| anti-eIF4A3 (rabbit mAb) | abcam | ab180573 | 1:250 |
| anti-eIF4E (rabbit mAb) | acam | ab33766 | 1:250 |
| anti-eIF4G1 (rabbit pAb) | Cell signalling | 2858 | 1:200 |
| anti-eIF4H (D85F2) XP® (rabbit mAb) | Cell signalling | 3469 | 1:400 |
| anti-PABP (rabbit pAb) | abcam | ab21060 | 1:400 |
| anti-FUS/TLS (rabbit mAb) | Cell signalling | 67840 | 1:800 |
| anti-TAF15 (8TA-2B10) (mouse mAb) | Invitrogen | MA3-078 | 1:250 |
| anti-PTBP1 (rabbit pAb) | Proteintech | 12582-1-AP | 1:100 |
| anti-eIF2α (rabbit mAb) | Santa Cruz<br>Biotechnology | sc-133132 | 1:500 |
| anti-phospho-eIF2α (S51) (mouse mAb) | abcam | ab32157 | 1:5'000 |
| anti-α-tubulin | Proteintech | 66031-1-Ig | 1:20'000 |
| Streptavidin Alexa 594 conjugate | Molecular Probes | S-32356 | 1:400 |
| Avidin, NeutrAvidin HRP conjugate | invitrogen | A2664 | 1:2'500 |
| donkey anti-rabbit Alexa 594 | Jackson<br>ImmunoResearch | 711-585-152 | 1:400 |
| donkey anti-mouse 647 | Jackson<br>ImmunoResearch | 715-605-150 | 1:400 |
| Peroxidase-conjugated Donkey anti-Mouse | Jackson<br>ImmunoResearch | 715-035-151 | 1:5'000 |
| Peroxidase-conjugated Donkey anti-Rabbit | Jackson<br>ImmunoResearch | 711-035-152 | 1:5'000 |

Image acquisition by fluorescence microscopy was done on a DeltaVision Elite High-Resolution widefield imaging system (General Electric) or a Nikon Ti-2 cicero spinning disk confocal microscope (Nikon) using 60x or 100x oil immersion objectives. For each imaged point a Z-stack with a Z resolution of 0.2um was acquired. Images acquired with the DeltaVision widefield system were additionally deconvolved with the in-built software softWoRx. The post-processing including a maximum intensity Z projection was done with the software Fiji and figures were assembled using the FigureJ plug-in (39,40). Quantification of signal intensity in cytoplasma was done by segmenting cytoplasma based on G3-A-G channel using Ilastik (41). Intensity of second channel in segmented cytoplasma was measured in Fiji and the average intensity on a per image base (normalized with area) was plotted in RStudio. One-way ANOVA and subsequent pairwise t-test was performed to determine significance.

### Live-cell imaging and image analysis

17Cl1:G3AG cells were seeded in 8-chambered glass bottom slides (ibidi, 80807) and infected with SFV or MHV at an MOI 10. Infection was synchronized by first incubating the slide on ice for 1h, followed by incubation at 37°C for 1.5h. Infection medium was exchanged with fresh MEM+/+/+ supplemented with 1:40 HEPES (gibco). Medium for MHV infected samples additionally contained 1uM HR2 peptide to prevent syncytia formation. Cells were imaged on a DeltaVision elite system fluorescence microscope equipped with a 40x oil immersion objective, an on-stage incubator chamber at 37°C, and 5% CO2 supply. Multiple points per condition were imaged from 2hpi up to 24hpi with an imaging interval of 10min.

Data analysis wase done with an image post-processing and quantification workflow based on Fiji and Ilastik. In short, an initial processing was done in Fiji including a maximum intensity Z projection. Next, Ilastik was used to train a pixel classification model on a subset of imaging data to differentiate between stress granules and background. The trained model was subsequently used to generate pixel prediction masks for all image series, which were further used to segment and count SGs in each image with an automated script in Fiji. The cell number for each image series was derived by manually counting the cells in selected frames throughout the imaging period. The number of SGs for each image was normalized with the corresponding cell number. The mean value over multiple image series was calculated for each condition and plotted using RStudio and the packages tidyverse and ggplot2 (42).

### SILAC labeling with isotopically labelled amino acids

All SILAC experiments were carried out with special medium either containing heavy isotope labelled arginine/lysine or normal isotope labeled arginine/lysine. MEM without L-arginine and L-lysine (MEM for SILAC, Thermo Fisher) was supplemented with 10% (v/v) FBS, 100μg/ml streptomycin, 100 IU/ml penicillin (Gibco), 1x NEAA (Gibco) and 0.25 mM L-proline:HCl (Thermo Fisher). The “light medium” was additionally supplemented with 0.6 mM normal isotope labelled L-arginine:HCl (Thermo Fisher) and 0.4mM normal isotope labelled L-lysine:2HCl (Thermo Fisher). The “heavy” medium was additionally supplemented with heavy isotope labelled L-arginine:HCl (^13^C_6_, ^15^N_4_; Cambridge Isotope laboratories) and heavy isotope labelled L-lysine:2HCl (^13^C_6_, ^15^N_2_; Cambridge Isotope laboratories).

Two batches of recombinant 17Cl1:G3-A-G cells were propagated on either heavy or light medium for a minimum of three weeks, and incorporation of isotopically labelled amino acids was verified by mass spectrometry.

### APEX2-mediated proximity biotinylation

Recombinant 17Cl1:G3-A-G cells propagated on either heavy or light SILAC medium were seeded in 10cm dishes two days prior to infection. Dishes for 2^nd^ and 3^rd^ SILAC experiments further contained a glass coverslip for visual verification of proximity labelling by IF (staining procedure described above). Cells were infected with either MHV-A59 at MOI 5 (1^st^ experiment) or 10 (2^nd^ experiment), or SFV at MOI 10. Infection media were additionally supplemented with 25 mM HEPES (Gibco). Samples for arsenite treatment or mock infection were incubated in fresh SILAC medium. The infection was synchronized by incubation of the dishes on ice for 1h, followed by 1.5h at 37°C. Infection medium was aspirated, cells were washed 3x with PBS, and fresh prewarmed SILAC medium was added. 30min prior to the respective labelling time point medium was replaced by fresh prewarmed SILAC medium supplemented with 0.5mM (1^st^ experiment) or 2.5mM (2^nd^/3^rd^ experiment) biotin phenol (BP). Samples for negative labelling control received medium without BP. APEX2-mediated proximity labelling was carried out by direct addition of hydrogen peroxide to the medium at a final concentration of ∼1 mM. After incubation of 60sec at room temperature the medium was removed, and cells were washed 3x with quencher solution consisting of PBS supplemented with Trolox (5 mM, Sigma), sodium ascorbate (10 mM, Sigma), and sodium azide (10 mM, Sigma). The quencher solution was freshly prepared for each labelling time point. Cells were lysed in 1ml lysis buffer (50 mM Tris-HCl pH 7.4, 500mM NaCl, 0.2% SDS, 1x cOmplete Mini Protease inhibitor cocktail (Roche), 5 mM Trolox, 10 mM sodium ascorbate, and 10 mM sodium azide in H_2_O), scraped off and collected in a tube. Triton-X100 was further added to the lysates to a final concentration of 2%. Next, the lysates were homogenized using a sonicator or MagNA lyser tubes (Roche), and cleared by centrifugation at 12’000 rpm for 10 min at 4°C. The protein concentration of each lysate was assessed by 660nm protein assay (Pierce), and light and corresponding heavy samples were mixed in a 1:1 protein ratio, 500ug of each. For the affinity purification, 100ul (1^st^ experiment) or 500ul (2^nd^/3^rd^ experiment) streptavidin coated magnetic beads (Pierce™ Streptavidin Magnetic Beads, Thermo Fisher) for each mixed lysate were washed 2x with lysis buffer. The mixed lysates were then incubated on the beads overnight at 4°C on a rotator. The next day, the supernatants were collected and stored for further analysis. A stringent washing procedure of the beads was performed consisting of two washes with buffer 1 (2% SDS in H2O), one wash with wash buffer 2 (2M Urea, 10mM Tris-HCl pH 7.4), two washes with wash buffer 3 (0.1% (w/v) deoxycholate, 1% (vol/vol) Triton X-100, 1mM EDTA, 500mM NaCl, 50mM HEPES pH 7.5 in H2O), two washes with wash buffer 4 (0.5% (w/v) deoxycholate, 0.5% (vol/vol) NP-40, 1mM EDTA, 250mM LiCl, 10mM Tris-HCl pH 7.4 in H2O) and a final wash with 50mM Tris-HCl pH 7.4. Each wash included 8min incubation on a rotator, followed by 1min incubation on a magnetic rack. To elute the proximity labelled proteins, the beads were incubated in 50ul elution buffer at 95°C for 10min on a shaking heat block (600rpm). The eluates were stored at -80°C until mass spectrometric analysis. The proximity labelling and affinity purification was verified by Western blot analysis of lysates, eluates and flowthroughs using horse radish peroxidase (HRP) conjugated streptavidin (Dako) or NeutrAvidin (Invitrogen) to detect biotinylated proteins. For the 2^nd^ and 3^rd^ experiment the subcellular localization of biotinylated proteins was further assessed by immunofluorescence microscopy using AlexaFluor 594 conjugated streptavidin (Invitrogen)(procedure described above).

### Mass spectrometry analysis

After affinity purification, eluates were fractionated according to protein size by SDS-PAGE and subsequent cutting of gel into 1mm3 cubes. The samples were digested in-gel and subjected to liquid chromatography tandem mass spectrometry (LC-MS/MS). In short, the gel cubes were reduced, alkylated and digested by trypsin as described previously (43). After loading, peptides were eluted in back flush mode onto the analytical self-packed nano-column (Acquity^TM^ CSH C18, 1.7µm, 100A, 0.075 mm i.d. x 200mm length, Waters, Baden, Switzerland) using an acetonitrile gradient of 5% to 40% solvent B (0.1% Formic Acid in water/acetonitrile 4,9:95) in 60 min at a flow rate of 350nL/min. The column effluent was directly coupled to a Fusion LUMOS mass spectrometer (Thermo Fischer, Bremen; Germany) via a nano-spray ESI source.

Data acquisition was made in data dependent mode with precursor ion scans recorded in the orbitrap with resolution of 120’000 (at m/z=250) parallel to top speed fragment spectra of the most intense precursor ions in the Linear trap for a cycle time of 3 seconds maximum.

Mass spectrometry data was analyzed as previously described (44). Briefly, proteins were identified by MaxQuant (version 2.5.1.0) against mouse protein database (SwissProt, release April 2025_01) and MHV strain A59 database (UniprotKB, release March 2023) (45). The initial precursor mass tolerance was set to 10 ppm, and that of the fragment peaks to 0.4 Da. Enzyme specificity was set to strict trypsin, and a maximum of three missed cleavages were allowed. Carbamidomethylation on cysteine was set as a fixed modification, while methionine oxidation, and protein N-terminal acetylation were set as variable modifications.

After removal of common contaminants and inversing reversed ratios, 1-sample t-tests were performed on the MaxQuant normalized ratios, provided the ratios were measured in all replicates. Adjusted p-values for multiple testing were corrected by the R fdrtool package (46).

Protein intensities were furthermore calculated as the median of the log_2_-transform of the intensities of all peptides allocated to the protein, using the heavy and light channels as separate samples, as previously described (12). Normalization was performed by the global median method, whereby the viral proteins as well as any previously known SG proteins (Curated list by Markmiller and colleagues (12)) were excluded from the calculation of the normalization coefficients. Missing protein intensities were imputed in the following manner, provided there was at least two identifications in at least one group: if there was at most two identifications in the group of replicates, the missing values were drawn from a Gaussian distribution of width 0.3 centered at the sample distribution mean minus 2.5x the sample standard deviation, otherwise the Maximum Likelihood Estimation (MLE) method was used (47). Log_2_ fold changes (Log2FC) between given conditions were calculated by using mean Log2 of the respective conditions. Differential expression tests were performed by applying an Empirical Bayes test (48), and the FDR-adjusted p-values (Benjamini and Hochberg multiple test correction (49)) were calculated. Significantly enriched or depleted proteins (adj. p-value ≤ 0.05 for large Log2FC, and |Log2(FC)| > 2 for individual comparisons or > log2(2.5) for SILAC based comparisons) were determined between different conditions. Data visualization was performed in R: Gene Ontology analysis was performed using the clusterProfiler (50). Redundant GO terms for each condition were reduced based on semantic similarity (similarity threshold: 0.7, method: relevance) using rrvigo package (51). Venn diagrams were generated with VennDiagram package (52). Further data analysis and visualization were performed using ggplot2 and tidyverse packages.

### Western blot analysis

17Cl1:G3AG cells were grown in 12-well lates and infected with MHV or SFV at an MOI of 10, or mock infected as described above. At the indicated time points, medium was removed, cells were washed once with PBS and lysed in lysis buffer (M-PER™ Mammalian Protein Extraction Reagent, Thermo Scientific) supplemented with cOmplete mini protease inhibitor (Roche) and phosphatase inhibitor (Thermo Scientific). Protein concentration was determined using Pierce BCA Potein assay (Thermo Scientific) and 15ng total protein was mixed with Lämmli buffer (supplemented with β-mercaptoethanol) and incubated at 95°C for 5min. Samples were run on an SDS-PAGE (4-20% gradient, SurePAGE GenScript) and blotted onto nitrocellulose membranes. Membranes were blocked in TBS + 0.1% Tween-20 (TBS-T) supplemented with 5% milk powder for 1h at room temperature. Staining with primary (anti-eIF2α, anti-phospho-eIF2α, α-tubulin) and secondary antibodies was done in TBS-T + 0.5% milk over night at 4°C or for 1h at room temperature respectively. Used antibody dilutions are found in Tab 1. Membranes were washed 3x with TBS for 5min. Membranes were developed with WesternBright ECL HRP substrate (Advansta) and imaged on a FusionFX. Band intensities were measured in Fiji and plotted in RStudio. In short, intensity for eIF2α and p-eIF2α bands were loading corrected with the corresponding loading control signal. The p-eIFα/eIF2α ratio was calculated and normalized with the Arsenite sample.

### smRNA FISH

17Cl1:G3AG cells were seeded on glass coverslips and infected with MHV or SFV at an MOI of 0.1 as described above. The smFISH protocol used to visualize viral mRNAs in this study was executed as previously detailed by Dave et al. (2023) (53). Cells were fixed at the specified time points post-infection using 4% paraformaldehyde in PBS for 20 minutes. The cells were then permeabilized with 0.5% Triton X-100 solution for 10 minutes and pre-blocked for 15 minutes in wash buffer consisting of 2xSSC (Invitrogen) and 10% v/v formamide (Ambion). Hybridization with fluorescently labeled oligos was then carried out using a hybridization buffer containing 150 nM smFISH probes specific to MHV genomic RNA (nsP3 gene), S-gene, SFV genomic RNA (nsP2 gene), or mmATF4 mRNA, along with 2xSSC, 10% v/v formamide, and 10% w/v dextran sulfate (Sigma), for 4 hours at 37°C. Following hybridization, cells were washed twice with wash buffer for 30 minutes each and subsequently 3x with PBS. Coverslips were mounted on slides with ProLong Gold antifade reagent containing DAPI (Molecular Probes). The 20-nucleotide oligonucleotides targeting various viral RNAs were designed using the Stellaris probe designer and conjugated to either Atto-565 or Atto-633, as described in the protocol by Gasper et al. (2017) (54).

Images were acquired using a Zeiss AxioObserver7 inverted microscope equipped with a Yokogawa CSU W1-T2 spinning disk confocal scanning unit, a Plan-APOCHROMAT 100x 1.4 NA oil objective, an sCMOS camera, and an X-Cite 120 EXFO metal halide light source. Imaging was performed with 21 z-stacks at 0.24 µm intervals.

Microscopy images were analyzed for colocalization in Fiji. The images were cropped to include a single cell per image and intensity of signal across multiple channels were measured using the plot profile function in ImageJ across the ROI (straight line across the object of interest). The plot profiles across channels were then plotted in python using seaborn.

## Acknowledgments

We want to thank Prof. Dr. Oliver Mühlemann from the Department of Chemistry, Biochemistry and Pharmaceutical Sciences at the University of Bern for providing us with the Semliki Forest virus strain 4.

The mass spectrometry experiments were conducted at the Proteomics and Mass Spectrometry Core Facility, Department of BioMedical Research (DBMR) of the University of Berne, Switzerland.

Microscopy was performed on instruments supported by the Microscopy Imaging Center (MIC), University of Bern, Switzerland.

This study was supported by the Swiss National Science Foundation SNSF (NCCR RNA & Disease, grant number 51NF40-205601.

## Supplementary information

**S1 Fig. Infection time course to assess SG formation during MHV infection.** 17Cl1 mouse fibroblast cells were infected with MHV-A59 at an MOI of 10. Cells were fixed at indicated time points and subjected to indirect immunofluorescence staining targeting the main SG marker G3BP1 (green) and dsRNA (red) as an intermediate of viral replication.

**S2 Fig. Experimental setup for SG proteome comparisons based on proximity labelling and quantitative proteomics using a stable isotope labelling in cell culture (SILAC) approach.**

(A) For SILAC based comparisons of two conditions, two batches of 17Cl1:G3-A-G cells are propagated on medium either containing heavy isotope labelled L-arginine/L-lysine or unlabeled (“light”) L-arginine/L-lysine. Cells are seeded in 10cm dishes. The treatments to compare are done, one on heavy cells and the other on light cells. APEX2-mediated proximity labelling is performed for both samples individually. After lysis, heavy and light lysates are mixed in a 1:1 protein ratio and a subsequent affinity purification of biotinylated proteins is performed on the mixture. Biotinylated proteins are identified by quantitative proteomics and a ratio of abundance between the compared treatments is calculated for each protein based on the intensity of heavy and light forms. (B) Five SILAC comparisons were performed to compare the SG proteome between oxidative stress (sodium arsenite treatment) and MHV infection. Comparison of each stressor with mock (comparison 1 and 4), comparison of each stressor with and without proximity labelling (comparison 2 and 5), and direct comparison of both stressors (comparison 3).

**S3 Fig. Comparison of different stress stimuli with and without APEX2-mediated proximity labelling.**

(A) Volcano plot shows the mean log2 fold change of proteins between sodium arsenite treated cells (oxidative stress) with (Ars APX+) and without (Ars APX-) APEX2-mediated proximity labelling of SG proteome plotted against the –log10 adjusted p value. Cells were treated with 250uM NaAsO2 for 30min. Proximity labelling was done with or without addition of biotin phenol. (B) Volcano plot shows the mean log2 fold change of proteins between MHV infected cells (oxidative stress) with (MHV APX+) and without (MHV APX-) APEX2-mediated proximity labelling of SG proteome plotted against the –log10 adjusted p value. Cells were infected with MHV-A59 (MOI 5). Proximity labelling was done at 8hpi with or without addition of biotin phenol. Proteins from our SG associated proteins reference list are indicated in orange, main SG marker proteins are labeled and indicated in green, and viral proteins are labeled and indicated in red. The selected significance levels for log2 fold change (>=2 and <=-2) and adjusted p value (<0.05) are shown with dashed lines.

**S4 Fig. Validation of APEX2-mediated proximity labelling and affinity purification in samples for mass spectrometry.**

(A) Immunofluorescence staining for biotinylated proteins (magenta) and dsRNA (red) of infected and proximity labelled samples for proteomics analysis. Heavy or light isotope labelled 17Cl1:G3-A-G cells (green) were grown in 10cm dishes each containing one glass coverslip. Cells were infected with MHV or SFV at MOI 10, mock infected or treated with sodium arsenate. APEX2-mediated proximity labelling was performed at the indicated time points. After quenching the cells on the coverslips were fixed and stained.

Images were acquired with a widefield fluorescence microscope and processed in Fiji. Scale bar: 20um. Presence of biotinylated (proximity labelled) proteins in (B) the individual lysates, and (C) the eluates and flowthroughs after affinity purification on streptavidin-coated magnetic beads was assessed by Western blot analysis. Membranes were stained with HRP-conjugated NeutrAvidin.

**S5 Fig. Differential protein abundance analysis of the G3-A-G microenvironment between MHV- and SFV-infected cells and mock controls based on proximity labelling and quantitative proteomics.**

(A) Differential abundance of proteins between unstressed, proximity labeled (mock APX+) and unstressed, unlabeled (mock APX-) cells. Volcano plot displays the mean log2 fold change based on the heavy/light protein ratio between the conditions (n = 3) plotted against the -log10 adjusted p value. Proteins contained in the reference list for SG associated proteins are indicated in orange, the major SG markers are labeled and indicated in green. Dashed lines show the significance criteria for log2 fold change >log2(2.5) and <- log2(2.5) respectively, and adjusted p value < 0.05. (B) Log2 fold change density distribution of total identified proteins (gray bar plot), and the corresponding subset of previously with SG associated proteins based on a reference list (orange curve). Results for direct comparison of MHV and SFV infections at pre SG time point (top), peak SG time point (middle) and post SG time point (bottom), based on volcano plots in fig. 5. (C) Venn diagram visualizing overlap between significantly enriched protein fractions (red: mock APX+ / blue: mock APX-), all identified proteins of this comparison (gray) and reference list of SG associated proteins (orange) for the unstressed proximity labeling control (mock). (D) Venn diagram visualizing overlap between significantly enriched protein fractions (red: MHV / blue: SFV), all identified proteins (gray) and reference list of SG associated proteins (orange) for the time point prior to SG formation (pre SG).

**S6 Fig. Dissection of SG proteome induced by SFV, MHV or sodium arsenite using APEX2-mediated proximity labelling and quantitative proteomics.**

Schemes visualize the different stress stimuli compared to unstressed cells. (A) Different abundance of proteins between SFV-infected (5hpi) and uninfected (mock), proximity labelled cells. (B) Different abundance of proteins between MHV-infected (8hpi) and uninfected (mock), proximity labelled cells. (C) Different abundance of proteins between sodium arsenite treated and untreated (mock), proximity labelled cells. Volcano plots display the mean log2 fold change based on the heavy/light protein ratio between the conditions (n = 3) plotted against the -log10 adjusted p value. Proteins contained in the reference list for SG associated proteins are indicated in orange, the major SG markers are labeled and indicated in green. Dashed lines show the significance criteria for |log2FC| >log2(2.5), and adjusted p value < 0.05. For the MHV comparison the adjusted p value < 0.1 is additionally indicated (red dashed).

**S7 Fig. Significantly enriched proteins in SG microenvironment during different stress stimuli.**

Venn diagrams visualizing overlap between significantly enriched protein fractions for the stress condition, all identified proteins in the respective comparison (gray), reference list of SG associated proteins (orange) and proximity labelling independent hits (proteins detected but not significantly enriched in “stressor with proximity labelling” vs. “stressor without proximity labelling” comparisons, gray) for (A) SFV, (B) MHV (less stringent p value criteria of <0.1 used) and (C) sodium arsenite. For selected sections, the corresponding lists of proteins are provided. Proteins in bold are found for all three stressors, colored proteins are only found for one virus (SFV: blue / MHV: red). (D) GO term enrichment analysis for significantly enriched proteins for each stressor using ClusterProfiler and rrvigo. Displayed are the top 5 most significant terms found in each subset of proteins for biological processes. Circle size indicates number of proteins (counts) identified for GO term.

**S8 Fig. Comparison of G3-A-G microenvironment between different stress conditions and time points based on proximity labelling and quantitative proteomics.**

Volcano plots show the mean log2 fold change of proteins based on the individual conditions plotted against the –log10 adjusted p value. Proteins which have not been detected in one condition were imputed to receive a ratio between the compared conditions. Proteins from our SG associated proteins reference list are indicated in orange, main SG marker proteins are labeled and indicated in green, and viral proteins are labeled and indicated in red. Shapes indicate if protein was absent in one condition, while present in all replicates (n = 3) of the second condition (strict on/off), or at least present in 2 replicates of the second condition (on/off). The selected significance levels for |log2FC| ≥2 and adjusted p value (<0.05) are shown with dashed lines.

**S9 Fig. Subcellular localization of selected proteins in relation to SGs induced by oxidative stress, MHV or SFV infection.**

17Cl1:G3-A-G (green) were either infected with MHV or SFV at an MOI of 10 or treated with 250uM sodium arsenite for 30min to induce SGs. Unstressed cells (mock) were included as negative control. MHV infected samples were additionally supplemented with HR2 peptide to prevent syncytia formation. Cells were fixed at indicated time points after infection/treatment and subjected to an indirect immunofluorescence staining with antibodies targeting the main SG markers (A) G3BP1, (B) TIA1, (C) UBAP2L, and additional protein (D) eIF4A3 (red channel). Additional staining for double stranded RNA (dsRNA, grays), an intermediate of virus replication, was done to visualize virus infection. Cell nuclei were stained with DAPI (blue). Scale bars: 20um

**S10 Fig. Subcellular localization of FUS over time in SFV infected cells.**

(A) 17Cl1:G3-A-G cells (green) were infected with SFV at an MOI 10, mock infected or treated with sodium arsenite. Cells were fixed at indicated time points; arsenite sample after 30min, mock sample at 7hpi. Indirect immunofluorescence staining was performed targeting FUS (red) and dsRNA (grays). Scale bar: 20um. (B) Quantification of average fluorescence intensity of FUS signal on a per-image base. The cytoplasm was segmented based on the G3-A-G channel and the area and integrated density was measured. For each, the mean average intensity based on three images is plotted for each condition. The dots represent the average intensity for the individual images; the error bar represents the standard deviation. P-value of one-way ANOVA over all conditions indicated. A paired t test was performed between the mock condition and each infected/treated condition. *: p < 0.05; **: p < 0.01; ns: non-significant.

**S11 Fig. Subcellular localization of PTBP1 over time in SFV infected cells.**

(A) 17Cl1:G3-A-G cells (green) were infected with SFV at an MOI 10, mock infected or treated with sodium arsenite. Cells were fixed at indicated time points; arsenite sample after 30min, mock sample at 7hpi. Indirect immunofluorescence staining was performed targeting PTBP1 (red) and dsRNA (grays). Scale bar: 20um. (B) Quantification of average fluorescence intensity of PTBP1 signal on a per-image base. The cytoplasm was segmented based on the G3-A-G channel and the area and integrated density was measured. For each, the mean average intensity based on three images is plotted for each condition. The dots represent the average intensity for the individual images; the error bar represents the standard deviation. P-value of one-way ANOVA over all conditions indicated. A paired t test was performed between the mock condition and each infected/treated condition. *: p < 0.05; **: p < 0.01; ns: non-significant.

**S12 Fig. Subcellular localization of TAF15 over time in SFV infected cells.**

(A) 17Cl1:G3-A-G cells (green) were infected with SFV at an MOI 10, mock infected or treated with sodium arsenite. Cells were fixed at indicated time points; arsenite sample after 30min, mock sample at 7hpi. Indirect immunofluorescence staining was performed targeting TAF15 (red) and dsRNA (grays). Scale bar: 20um. (B) Quantification of average fluorescence intensity of TAF15 signal on a per-image base. The cytoplasm was segmented based on the G3-A-G channel and the area and integrated density was measured. For each, the mean average intensity based on three images is plotted for each condition. The dots represent the average intensity for the individual images; the error bar represents the standard deviation. P-value of one-way ANOVA over all conditions indicated. A paired t test was performed between the mock condition and each infected/treated condition. *: p < 0.05; **: p < 0.01; ns: non-significant.

**S13 Fig. Subcellular localization of mATF4 mRNA as representative host mRNA in relation to SGs induced during MHV infection using smRNA FISH.**

17Cl1:G3-A-G cells (blue) were infected with MHV at an MOI of 0.1. Medium was then supplemented with HR2 peptide to prevent syncytia formation. Cells were fixed at indicated time points and subjected to smRNA FISH with a probe targeting the mouse activating transcription factor 4 mRNA (mmATF4, green), and a probe targeting MHV genomic RNA (vRNA, magenta). Line plots display measured intensity of G3-A-G, mmATF4 and vRNA channels along the selected line (white) indicated in the inset. Cell nuclei were stained with DAPI (grays, only in merge). Scale bar: 5um

**S1 Movie SG formation during MHV and SFV infections monitored assessed by live-cell imaging.**

17Cl1 cells expressing a G3BP1-APEX2-GFP fusion protein (green) were infected with MHV or SFV at an MOI of 10, or mock infected. MHV infected samples were additionally supplemented with HR2 peptide to prevent syncytia formation. Cells were imaged every 10min on a DeltaVision widefield fluorescence microscope equipped with an on-stage incubator and 5% CO2 supply. Scale bar: 20um

**S1 Table Reference list of SG associated proteins related to Figures 2, 3, 5, and supplementary figures 3, 5, 6, 7, 8.** Proteins are derived from Markmiller et al. 2018 or the Mouse Genome Informatics (MGI) data base.

**S2 Table MaxQuant output for first APEX2-based SG proteome analysis comparing MHV and sodium arsenite, related to Figures 2 and 3, and supplementary figure 3.** The file contains protein intensities including corresponding analysis based on individual conditions (proteinGroups DE test) and heavy/light ratio (proteinGroups cleaned), an experimental design overview, and additional information.

**S3 Table MaxQuant output for second APEX2-based SG proteome analysis comparing MHV and SFV, related to Figures 5 and 6 and supplementary figures 5 and 8.** The file contains protein intensities including corresponding analysis based on heavy/light ratios (proteinGroups Normalized ratios) and individual conditions (proteinGroups DE test), an experimental design overview, and additional information.

**S4 Table MaxQuant output for second APEX2-based SG proteome analysis comparing MHV, SFV and arsenite against mock, related to supplementary figure 6 and 7.** The file contains protein intensities including corresponding analysis based on heavy/light ratios (proteinGroups Normalized ratios) and individual conditions (proteinGroups DE test), an experimental design overview, and additional information.

